# MEK signaling represents a viable therapeutic vulnerability of *KRAS*-driven somatic brain arteriovenous malformations

**DOI:** 10.1101/2024.05.15.594335

**Authors:** Carlos Perfecto Flores Suarez, Omar Ashraf Harb, Ariadna Robledo, Gabrielle Largoza, John J. Ahn, Emily K. Alley, Ting Wu, Surabi Veeraragavan, Samuel T. McClugage, Ionela Iacobas, Jason E. Fish, Peter T. Kan, Sean P. Marrelli, Joshua D. Wythe

## Abstract

Brain arteriovenous malformations (bAVMs) are direct connections between arteries and veins that remodel into a complex nidus susceptible to rupture and hemorrhage. Most sporadic bAVMs feature somatic activating mutations within *KRAS*, and endothelial-specific expression of the constitutively active variant KRAS^G12D^ models sporadic bAVM in mice. By leveraging 3D-based micro-CT imaging, we demonstrate that KRAS^G12D^-driven bAVMs arise in stereotypical anatomical locations within the murine brain, which coincide with high endogenous *Kras* expression. We extend these analyses to show that a distinct variant, KRAS^G12C^, also generates bAVMs in predictable locations. Analysis of 15,000 human patients revealed that, similar to murine models, bAVMs preferentially occur in distinct regions of the adult brain. Furthermore, bAVM location correlates with hemorrhagic frequency. Quantification of 3D imaging revealed that G12D and G12C alter vessel density, tortuosity, and diameter within the mouse brain. Notably, aged G12D mice feature increased lethality, as well as impaired cognition and motor function. Critically, we show that pharmacological blockade of the downstream kinase, MEK, after lesion formation ameliorates KRAS^G12D^-driven changes in the murine cerebrovasculature and may also impede bAVM progression in human pediatric patients. Collectively, these data show that distinct KRAS variants drive bAVMs in similar patterns and suggest MEK inhibition represents a non-surgical alternative therapy for sporadic bAVM.

## INTRODUCTION

<u>B</u>rain <u>a</u>rterio<u>v</u>enous <u>m</u>alformations (bAVMs) are abnormal direct connections between arteries and veins without an intervening capillary network. This arteriovenous shunt progressively remodels into a tangled and dilated nidus that is prone to leak or rupture(Lawton et al. 2015). Consequently, bAVMs are the main cause of hemorrhagic stroke in children and young adults and current approaches to bAVM management are continued observation or high-risk surgical intervention(Choi and Mohr 2005). However, until recently, the cause(s) of sporadic bAVMs were unknown. This gap in knowledge likely explains the lack of treatment options for bAVM. This changed after our landmark study identified somatic mutations in *KRAS* in the endothelium of most sporadic bAVM patients(Nikolaev et al. 2018).

*KRAS* encodes a small GTPase that can activate numerous downstream effector networks, such as the PI3K/AKT/mTOR pathway and the MEK/ERK kinase cascade(Simanshu et al. 2017). The *KRAS* variants we and others identified in bAVMs result in constitutive KRAS activation(Nikolaev et al. 2018; Goss et al. 2019; Hong et al. 2019; Oka et al. 2019; Bameri et al. 2021; Gao et al. 2022). Previously, we showed that mutant KRAS preferentially stimulates MEK/ERK signaling in cultured endothelial cells (ECs)(Nikolaev et al. 2018). Moreover, despite the fact that somatic *KRAS* variants occurred at a low variant allele frequency within the bAVM, histopathology revealed widespread ERK activation in patient bAVM samples(Nikolaev et al. 2018). In recent follow-up studies, we demonstrated that endothelial-specific gain of KRAS activity (i.e., G12D and G12V) is sufficient to induce bAVMs both in vitro and in vivo(Fish et al. 2020; Soon et al. 2022). Expression of mutant KRAS upregulated genes involved in angiogenesis in a MEK-dependent manner in cultured ECs, while MEK inhibition rescued arteriovenous shunts in a zebrafish model(Fish et al. 2020). A subsequent study showed that AAV-mediated delivery of the G12V *KRAS* variant also induced sporadic bAVM formation in mice(Park et al. 2021). Despite these findings, our understanding of how different KRAS variants remodel the murine cerebrovasculature, as well as their impact on vessels throughout the central nervous system, remains unclear. Furthermore, how these variants impact murine survival, as well as cognitive function, and how AAV and genetic models compare to one another is unknown.

Encouragingly, studies in zebrafish (KRAS^G12V^)(Fish et al. 2020), and AAV-KRAS^G12V^ transduced mice, showed that bAVMs were prevented by treatment with the MEK inhibitor Trametinib (Park et al. 2021). Similarly, a later study showed that Trametinib alleviates endothelial dysplasia in the neonatal cerebrovasculature in a *Kras^G12D^* mouse model, although both the AAV and later mouse study failed to examine the impact on established lesions(Nguyen et al. 2023).

Herein, using our previously validated murine genetic *Kras^G12D^*-model of sporadic bAVM, we leveraged 3D analysis through micro-CT imaging to define the frequency and location of these anomalies within the mouse brain over time. Notably, the frequency and anatomical location of bAVMs in this genetic G12D model differ significantly from a recently published study where mice were transduced with *AAV-KRAS^G12V(Park^ ^et^ ^al. 2021)^*. To determine if unique G12 variants drive bAVM in different locations, we developed a second, novel genetic model of sporadic bAVM. While KRAS^G12C^ mutations are rare in sporadic bAVMs (Priemer et al. 2019), there is hope that this mutation can be directly targeted with recently developed G12C-specific inhibitors (Canon et al. 2019). We show for the first time that G12C is sufficient to drive bAVM in mice, and these lesions occur with a frequency comparable to G12D, and in a similar region within the posterior cortex of the mouse brain. This contrasts with AAV-KRAS^G12V^ animals, which feature lesions in the olfactory bulb (Park et al. 2021). Next, micro-CT imaging and quantitative 3D morphometric analysis revealed these mutant vessels exhibit increased diameters and undergo extensive remodeling. Furthermore, we show that the presence of these bAVM lesions correlates with decreased motor function and cognitive ability in older mice. Notably, initiating pharmacological blockade of the downstream kinase, MEK, after lesion initiation decreased bAVM incidence and ameliorated vessel remodeling. To translate these results to the clinic, we show that MEK inhibitor treatment may slow bAVM progression in human pediatric patients. Overall, these data suggest mutant KRAS progressively remodels the cerebrovasculature with age, and that MEK activity represents a viable therapeutic target to reduce sporadic bAVM disease progression.

## RESULTS

### Postnatal *Kras^G12D^* expression in the murine CNS endothelium leads to bAVM formation preferentially in the posterior cortex

To characterize the morphological consequences of *Kras* gain of function in the murine cerebrovasculature, we employed our previously validated somatic mouse model (Fish et al. 2020) in which expression of a constitutively active mutant *Kras^G12D^* transcript depends upon the activity of a tamoxifen-inducible, brain EC-specific Cre recombinase driver line, *Slc1o1c1(BAC)-CreER* (Ridder et al. 2011). In this model, a LoxP flanked stop cassette is inserted downstream of the *Kras* promoter at the endogenous murine *Kras* locus, blocking transcription (Jackson et al. 2001). However, in the presence of Cre recombinase, the stop cassette is recombined and excised, juxtaposing the native *Kras* promoter and the exon containing the G12D mutation (**Fig. 1A**). Previously, we used this inducible <u>b</u>rain <u>e</u>ndothelial <u>c</u>ell *Kras^G12D^* gain of function line (hereafter referred to as *ibEC-Kras^G12D^*) to model somatic gain of KRAS activity in the postnatal central nervous system (CNS) endothelium(Fish et al. 2020). As in our previous studies, perfusion of fluorescent-labeled lectin revealed bAVMs in over half of all *ibEC-Kras^G12D^* mice at 2 months of age (Control n=0/17, Mutants n=11/16; **Fig. 1B, 1C**). Because visualization of the entire brain vasculature following lectin perfusion “en volume” requires tissue clearing and imaging by lightsheet fluorescent microscopy, as well as complicated post-imaging processing, we chose a streamlined alternative for generating 3D images of the cerebrovasculature, namely micro-computed tomography (micro-CT) (Hong et al. 2020).

**Figure 1.**
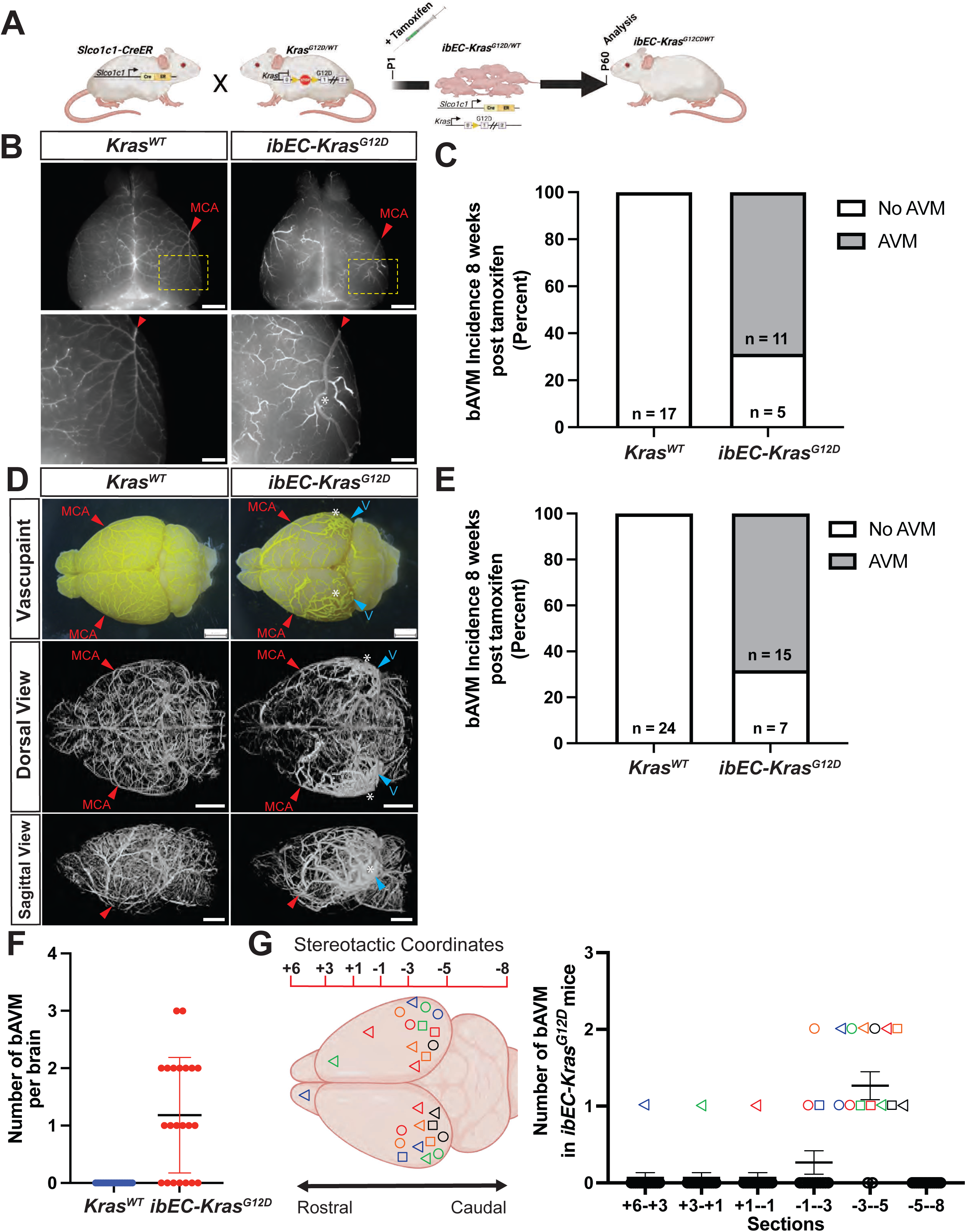
Endothelial-specific KRAS^G12D^ expression induces bAVMs preferentially within the caudal region of the mouse brain. (**A**) Breeding scheme for generating inducible <u>b</u>rain <u>e</u>ndothelial <u>c</u>ell *Kras^G12D^* mutant mice (*ibEC-Kras^G12D^*). Tamoxifen is delivered to pups at P1, and tissues are harvested at 8 weeks (postnatal day 60, P60) of age. (**B**) Panels in the top row are representative whole mount direct epifluorescence dorsal view images of the brain following vascular perfusion of fluorescent-conjugated tomato lectin. Yellow boxed areas are magnified and shown in the lower panels, with the red caret denoting the mid-cerebral artery [MCA]. The white asterisk highlights the presence of a brain arteriovenous malformation with a direct connection between an artery and a dilated draining vein in a 2-month-old *ibEC-Kras^G12D^* animal. (**C**) Quantification of bAVM incidence, as determined by lectin perfusion and epifluorescence microscopy, at 8 weeks of age after post tamoxifen induction at P1. (**D**) Top row shows representative phase microscopy images of the dorsal surface of the brain following arterial perfusion of vascupaint contrast agent, revealing normal arterial vessel patterning in *Kras^WT^* mice and obvious bAVMs in P60 *ibEC-Kras^G12D^* mice. The middle row shows representative dorsal views taken from 3D reconstructions of micro-CT imaged brains, while the bottom row shows sagittal views (olfactory bulbs to the left, cerebellum to the right) from the same representative brain. Note the extensive vessel remodeling in the *ibEC-Kras^G12D^* mice at the sites of two bAVMs indicated by the blue arrows. V=vein, MCA=mid cerebral artery, asterisk=bAVM. (**E**) Quantification of bAVM incidence at P60, as determined by perfusion with Vascupaint contrast agent and micro-CT imaging, after tamoxifen induction at P1. (**F**) The total number of bAVMs per mouse brain at two months of age. (**G**) Quantification and location of lesion incidence across Bregma stereotactic regional coordinates. Each unique color (blue, red, green, etc.) and shape combination (square, triangle, circle) correspond to the bAVMs identified in a single animal.

The perfusion reagent we employed, Vascupaint, serves as a vascular CT contrast agent that predominantly accumulates in the arterial vasculature. Though the agent passes through the capillaries, the properties of the material do not favor accumulation within the venous circulation. This feature of the contrast agent allows us to effectively restrict our imaging to the arterial vasculature. However, in 2-month-old *ibEC-Kras^G12D^* mutants, connections between the middle cerebral artery [MCA] and the transverse sinus vein (TSV) were observed, as well as additional connections between the MCA and the superior sagittal sinus (**Fig. 1D**). Quantification of these anomalous connections, or arteriovenous shunts at 2 months of age confirmed endothelial-specific recombination of the mutant *Kras^G12D^* allele robustly induces bAVMs (Control n=0/24, Mutants n=15/22; **Fig. 1E**). We would note that bAVM incidence was comparable between each method (68.8% for lectin vs 68.2% for micro-CT), although Vascupaint perfusion may facilitate bAVM identification due to the confinement of the contrast agent to the arterial network under normal circumstances.

Previously, we anecdotally noticed that *ibEC-Kras^G12D^* mice usually feature between one and three bAVMs per animal(Fish et al. 2020). Quantification confirmed this observation (**Fig. 1F**). Given the consistent induction of 1-2 bAVMs per animal, we wondered if there was any pattern to the location of these anomalies. Accordingly, micro-CT imaged brains were segmented in 2D along the rostral to caudal (anterior to posterior) axis using a stereotactic scale (Bregma +6 to Bregma -8, where Bregma is position 0; Bregma refers to a point in the skull where the coronal and sagittal sutures intersect)(Franklin and Paxinos 2013). Interestingly, the majority of bAVMs (∼93%) were observed in the -1 to -3 and -3 to -5 regions, respectively, although they were also rarely located in the caudal- (-5 to -8) (e.g. cerebellum) or rostral-most (+6 to +3) regions (e.g. olfactory bulbs) (**Fig. 1G**).

### Postnatal *Kras^G12C^* expression in the murine CNS endothelium leads to bAVMs predominantly in the posterior cortex

Given the identification of variants other than *KRAS^G12D^* in sporadic bAVM(Nikolaev et al. 2018; Priemer et al. 2019), we were curious if these other amino acid changes were sufficient to induce bAVMs, and if so, whether they yielded a similar or distinct pattern in terms of anomaly prevalence or location compared to G12D. Accordingly, we used mice harboring a *Kras^lox-stop-lox-G12C^* allele (Zafra et al. 2020) to generate inducible <u>b</u>rain <u>e</u>ndothelial <u>c</u>ell *Kras^G12C^* gain of function mice (hereafter referred to as *ibEC-Kras^G12C^*) (**Fig. 2A**). Delivery of tamoxifen at P1 did not impact survival to P60 and did not lead to frank intracranial hemorrhage (n=0/8) (**Fig. 2B,C**). However, perfusion with contrast agent, followed by micro-CT imaging, revealed that G12C robustly drove bAVM formation by 2 months of age (n=7/10) compared to control littermates (n=0/6), as connections between the middle cerebral artery [MCA] and the transverse sinus vein (TSV) were observed, as well as additional connections between the MCA and the superior sagittal sinus (**Fig. 2D,E**). Quantification revealed that G12C led to as many as 4 bAVMs per mouse, with a mean of 2 lesions per animal (**Fig. 2F**). The anomalies were most frequently observed in Bregma +3 to +1 and -3 to -5 (the latter region where most lesions arose in the G12D model), while no anomalies were detected in the cerebellum, and only 1 was found in the rostral-most olfactory bulb region (**Fig. 2G**).

**Figure 2.**
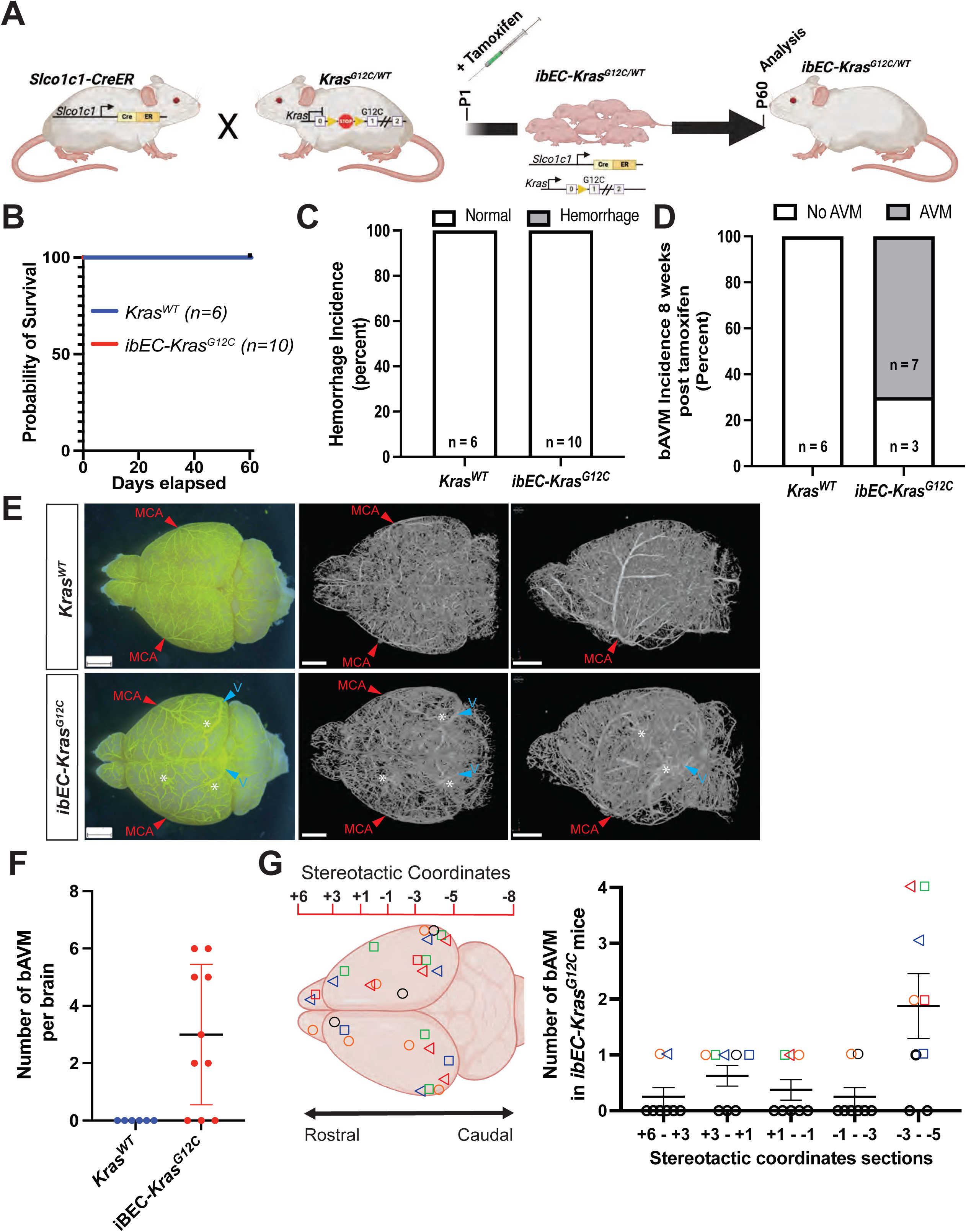
Endothelial KRAS^G12C^ expression is sufficient to induce bAVM by 2 months of age. (**A**) Mating scheme for generating inducible <u>b</u>rain <u>e</u>ndothelial <u>c</u>ell-specific *Kras^G12C^* mice (*ibEC-Kras^G12C^*). Pups are injected at P1 with tamoxifen and scored at 2 months of age for the presence of bAVM or hemorrhage. (**B**) Survival up to two months of age is indistinguishable between *ibEC-Kras^G12C^* mice and control littermates. (**C**) Intracerebral hemorrhage is not observed in either *ibEC-Kras^G12C^* mutants or control littermates. (**D**) Quantification of bAVM incidence at 2 months of age. (**E**) The top row shows representative images of a brain from a control, *Kras^WT^* animal, while the bottom row shows images of the brain from an *ibEC-Kras^G12C^* mouse. The far left column are phase microscopy images of the dorsal view of the brain (olfactory bulbs to the left, cerebellum to the right) following perfusion with vascupaint contrast agent. The middle and right column show representative reconstructions following micro-CT imaging, dorsal and sagittal views, respectively. scale bar = 2 mm. (**F**) Quantification bAVM number per animal (each dot represents one brain) in *Kras^WT^* (n=6) and *ibEC-Kras^G12C^* (n=8). (**G**) Graphical representation, and quantification, of location of lesion incidence across Bregma stereotactic regional coordinates. Each unique color (blue, red, green, etc.) and shape combination (square, triangle, circle) correspond to the bAVMs identified in a single animal.

### Postnatal mutant *Kras* expression in the murine CNS endothelium preferentially generates bAVM in the murine cortex

Given the evident regional preference for bAVM induction in the *ibEC-Kras^G12D^* and *ibEC-Kras^G12C^* models along the rostal-caudal axis, we wanted to determine if there was a structural preference for bAVM location in terms of the major sub regions of the grey matter (i.e. the cerebrum, cerebellum, and brain stem): specifically, the olfactory bulbs, cortex, hippocampus, striatum, thalamus, hypothalamus, and cerebellum. To determine if bAVM induction showed any regional preference in either of the G12D or G12C genetic models, we examined 2,150-micron thick virtual coronal sections (or “slabs”), moving along the rostral to caudal axis (olfactory bulbs to cerebellum). This analysis revealed that most *Kras^G12D^*-driven bAVMs (n=15 mice) occurred predominantly in the cerebral cortex (n=20/26), and infrequently within the olfactory bulb (n=1/26), hippocampus (n=4/26), and striatum (n=1/26) (**Fig. 3A, B**). For the *Kras^G12C^* model, most lesions (n=7 mice) were similarly confined to the cortex (n=22/29), followed by the hippocampus (n=3/29), olfactory bulbs (n=3/30), and hypothalamus (1/29) (**Fig. 3C, D**). In neither case were bAVMs observed in reconstructed sections from control littermates.

**Figure 3.**
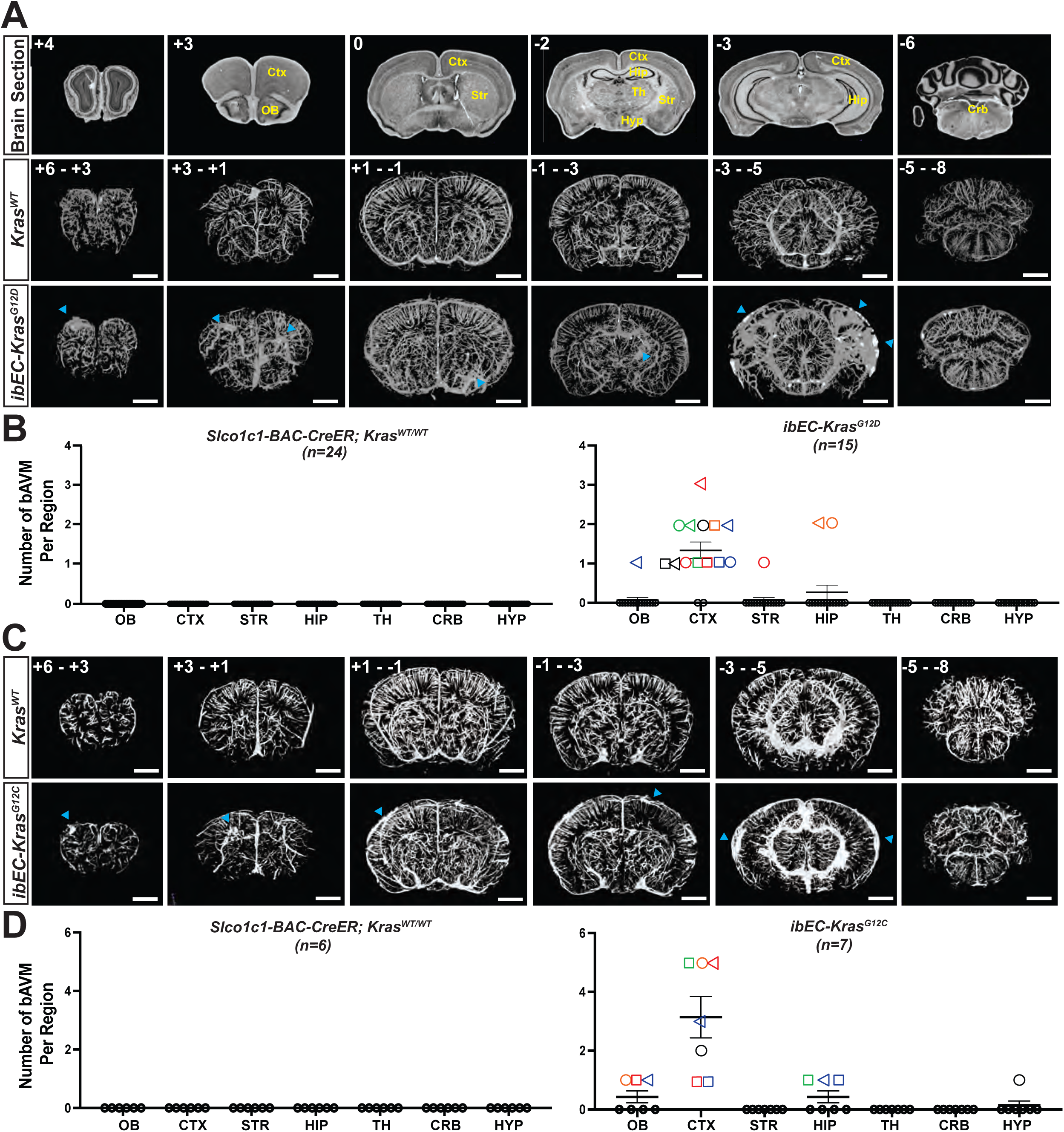
BAVMs in *ibEC-Kras^G12D^* and *ibEC-Kras^G12C^* mutant mice are predominantly located in the cortex. (**A)** The top row shows representative virtual sections of the adult mouse brain, coronal plane, progressing from rostral to caudal (left to right, respectively), with the plane of Bregma indicated in the upper left corner. The middle and bottom rows show representative 2D flattened slab reconstructions of approximately 2,150 microns obtained by micro-CT imaging following perfusion with Vascupaint contrast agent from 2-month-old control littermates and *ibEC-Kras^G12D^* mutants, respectively. Scale bar = 1000 microns. (**B**) Quantification of bAVMs by brain region in *Control* and *ibEC-Kras^G12D^* mutants at 2 months of age. (**C**) The top and bottom rows show representative 2D flattened slab reconstructions of approximately 2,150 microns obtained by micro-CT imaging following perfusion with Vascupaint contrast agent from 2-month-old adult *ibEC-Kras^G12C^* mutants and control littermates. Scale bar = 1000 microns. (**D**) Quantification of bAVMs by brain region in *Control* and *ibEC-Kras^G12C^* mutants at 2 months of age. OB=olfactory bulb; CTX=cortex; STR=striatum; HIP=hippocampus; TH=thalamus; CRB=cerebellum; HYP=hypothalamus.

Overall, the location and frequency of bAVMs in these two genetic models contrast with those where mice were transduced with an adeno-associated virus 2 (AAV2) variant reported to preferentially transduce the CNS endothelium known as AAV-BR1 (<u>AAV</u>-<u>Br</u>ain<u>1</u>)(Korbelin et al. 2016). In these AAV experiments, the synthetic CAG promoter drove *KRAS^G12V^* expression. Approximately 91% of *AAV2-BR1-KRAS^G12V^* mice developed a bAVM in the olfactory bulbs(Park et al. 2021), while in the *ibEC-Kras^G12D^* genetic model, 93% of bAVMs were observed in the posterior region of the brain, primarily in the cerebral cortex. Similarly, in our *ibEC-Kras^G12C^* model most bAVMs were located outside of the olfactory bulbs.

To determine if the spatial discrepancy between our studies and Park et al. was attributable to the promoter element used in their AAV model (as opposed to using the endogenous murine *Kras* promoter in the genetic G12D and G12C models described herein), we examined *AAV2-BR1-CAG* activity in the adult mouse. As reported previously, *AAV2-BR1-CAG-Cre* recombined a Cre reporter allele selectively in the brain (as opposed to other organs, such as the liver)(Korbelin et al. 2016). However, our data, in agreement with two recent reports(Santisteban et al. 2020; Krolak et al. 2022), show it primarily transduced a portion of the capillary endothelium but not large diameter vessels (**Supp. Fig. 1A,B**). Additionally, *AAV2-BR1-CAG-Cre* readily transduced cells outside of the CNS endothelium, as RFP signal was evident in neurons (**Supp. Fig. 1B**). Our data show that exchange of the CAG promoter for the lower-strength *EF1a* promoter partially mitigated this non-endothelial activity in the brain and retina (**Supp. Fig. 1C**).

In contrast, tamoxifen delivery at postnatal day 1 to *Slco1c1-CreER* mice led to robust recombination in the CNS endothelium in large and small diameter vessels, particularly within the cortex, as well as some non-endothelial cell types (**Supp. Fig. 2**). We previously showed *Slco1c1-CreER* induced uniform recombination within the brain across the rostral to caudal axis(Fish et al. 2020). However, while *Slco1c1-CreER* recombination of the floxed stop cassette preceding the open reading frame of mutant *Kras* is restricted to cell types in the CNS (in effect acting as a permissive signal), expression of either mutant G12D or G12C in our genetic knock-in models is driven by the endogenous *Kras* promoter.

Accordingly, we examined the spatial distribution of *Kras* transcripts in the adult murine brain using smFISH. Quantification revealed *Kras* transcripts appear to be unevenly distributed throughout the adult murine brain, while mRNA was significantly enriched in the endothelium of the cortex compared to the olfactory bulb and other regions of the brain (**Supp Fig. 3**), potentially explaining why these lesions preferentially arise in the posterior cortical region of *Kras* mutant mice. While other plausible mechanisms could potentially explain these disparate results, such as regional differences in hemodynamics and shear stress sensing, or heterogeneity in terms of cell signaling or even endothelial subtype composition across the brain(Dubrac et al. 2023), their direct role in KRAS-driven lesions remains to be determined. Regardless, given the regional preference for bAVM formation in the *ibEC-Kras^G12D^* and *ibEC-Kras^G12C^* mouse models, we wondered if bAVMs in human patients feature a similar spatial distribution.

### BAVMs in humans exhibit a regional preference in both location and hemorrhage frequency

To determine if bAVM lesions exhibit a stereotypical spatial distribution in human patients, Pubmed was queried to capture all manuscripts including the words “brain,” “arteriovenous malformation,” and “location,” limiting results to those written in, or translated to, English, yielding a total of 599 results, published between the year 1977 and 2023. 90 out of 599 screened articles met the inclusion criteria, as they describe patients with a diagnosis of non-familial bAVM, contained information on bAVM location, number of patients, and either age of patient at time of diagnosis or initial presentation of hemorrhage.

A total of 15,342 patients from 90 of these articles were categorized by anatomical location of bAVM. The most frequent location for bAVM was in the frontal lobe (n=3,438; 22.4%), followed by midline structures (n=2,822; 18.4%), and temporal lobe (n=2,337; 15.4%). The least common locations for bAVM were the cerebellopontine angle (n=18, <1%), pineal gland (n=13, <1%) and temporoparieto-occipital lobe (n=1, <1%) (**Fig. 4A-C**).

**Figure 4.**
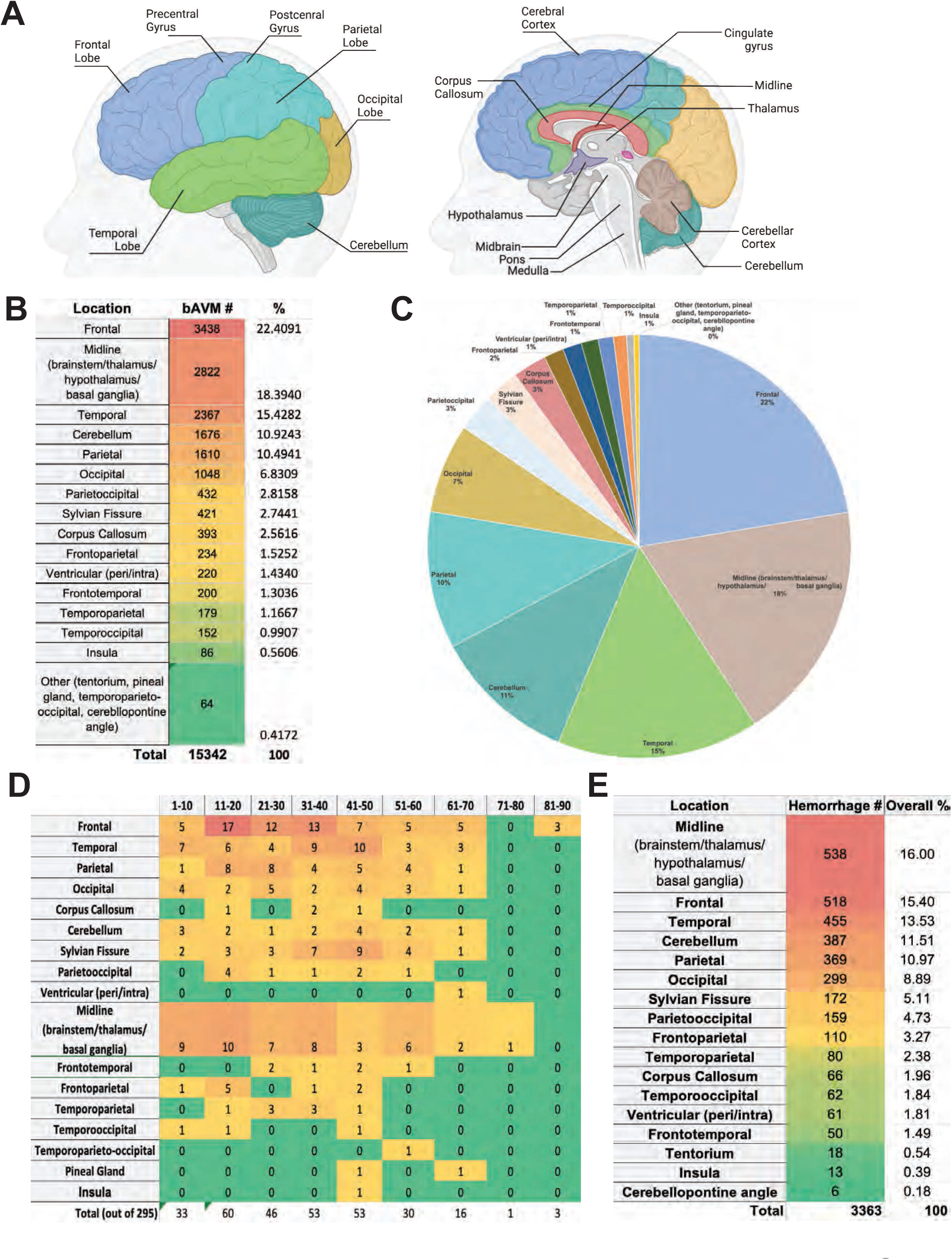
BAVMs in humans are located preferentially in the cortical lobes of the brain. (**A**) Representation of the regions of the human brain. (**B**) Heatmap showing the number and frequency of bAVM occurrence across various anatomical regions of the human brain from 15,342 patients. The most frequent location for bAVM was in the frontal lobe (n=3,438), followed by midline structures (n=2,822), and temporal lobe (n=2,367). The least common locations for bAVM were the tentorium, pineal gland, temporo-parieto-occipital and cerebellopontine angle (combined n=64 total cases). (**C**) Quantification of the bAVM frequency in these regions (shown as percentages of the total). (**D**) Categorizing the anatomical location of bAVM by age range. (**E**) Heatmap showing the incidence of intracranial hemorrhage for bAVM within each anatomical region, as well as the overall frequency of hemorrhage.

Forty-three of the reviewed articles contained information regarding the patient’s age for a total of 295 patients. Categorizing the anatomical location of bAVM by age range (**Fig. 4D**) revealed the most frequent location for bAVM in pediatric patients (age < 21 y/o) was the frontal lobe (n=22, age 1-20 y/o), followed by midline structures (n=19, age 1-20 y/o) and the temporal lobe (n=13, age 1-20 y/o). Middle-age adults (age >= 21 years old) showed a maximum frequency of bAVM in the frontal lobe (n=13, age 31-40 y/o) locations, while an older age range displayed a similar pattern with the most frequent location in temporal lobe (n=10, age 41–50 y/o) and midline structures (n = 6, age 51-60 y/o). No patients were recorded in the 71-80 y/o age range in this retrospective study and in the oldest cohort, all recorded cases were in the frontal lobe (n=3, 81-90 y/o) (**Fig. 4**).

Next, we analyzed this retrospective cohort to determine if the anatomical location of the sporadic lesion correlated with the incidence of intracranial hemorrhage. The anatomical location of bAVM and initial presentation of hemorrhage were compared from 3,363 patients from 27 reviewed articles (**Fig. 4E**). The most frequent anatomical location for bAVM hemorrhage was in midline structures (n=538, 16%), followed by frontal lobe (n=518, 16%), and temporal lobe (n=455, 13.53%). The least common locations for bAVM hemorrhage were found to be the cerebellopontine angle (n=6, <1%), precentral gyrus (n=13, <1%), and tentorium (n=18, <1%). Chi-square test results show that hemorrhage frequency was not evenly distributed and varied significantly between anatomical regions (p<0.0001), suggesting this frequency does not merely reflect the anatomical region where bAVM were most likely to occur.

Overall, these data suggest that, as in our murine model, sporadic bAVMs in human patients arise in stereotypical anatomic locations within the brain, and these data also indicate there may be a distinction where bAVMs are most frequently detected in pediatric versus adult patients. Notably, these analyses also show that the propensity for a bAVM to rupture varies by region, and hemorrhage incidence does not simply follow anatomical frequency. Collectively, these data suggest there may be unique features within the brain that predispose a region to AVM formation. Of note, the mutational status of *KRAS* in these lesions is unknown, and perhaps future analysis of case reports featuring endovascular biopsy and genetic diagnoses will determine if KRAS-induced lesions feature any anatomical preference or predisposition to intracranial hemorrhage.

### KRAS^G12D^ gain of function does not induce AVMs within the murine retinal vasculature or alter vascular patterning in the adult mouse retina

AVMs occur in the retina in murine models of HHT, a disease which can – albeit infrequently – feature bAVMs(Mahmoud et al. 2010; Tual-Chalot et al. 2014.; Ola et al. 2016; Crist et al. 2018). To determine the impact of mutant KRAS on the vasculature of the murine retina, *ibEC-Kras^G12D/+^* mice that were injected with tamoxifen at P1 were perfused with fluorescent lectin at 2 months of age to label the vasculature, and retinas were collected for immunohistochemical analysis and imaging. Antibody staining for alpha-smooth muscle actin (α-SMA), a marker of the smooth muscle cells that predominantly associate with mature arteries, failed to show any obvious arteriovenous shunts or malformations in the mutant or control adult retinas (n=7 and n=7, respectively) (**Supp. Fig. 4A, B**). Given the preservation of the normal alternating artery-vein patterning originating from the center of the optic nerve, and absence of arteriovenous shunts, we next asked whether the vessels in mutants featured any morphological differences compared to control animals.

Confocal imaging and analysis of the three layers of the retinal vasculature (superficial, intermediate, and deep plexus) (Rust et al. 2019)(**Supp. Fig. 4C**) failed to reveal any differences in vessel area, vessel density, vessel length, lacunarity and branching in *Kras* mutants compared to control littermates (n=4 for each genotype) (**Supp. Fig. 4D-G**). In agreement with a previous study, we also found that *Slco1c1-CreER* induced robust recombination in the arterial endothelium but largely spared the venous endothelium in the murine retina(Yang et al. 2020). However, we also detected strong CreER activity in the capillary endothelium (**Supp. Fig 5A,B**). Critically, the lack of observable AVMs in the adult retina using *Slco1c1-BAC-CreER* aligns with the results of Nguyen et. al., as they did not observe any defects in arteriovenous patterning at P8 using the pan-endothelial *Cdh5-PAC-CreER* driver(Nguyen et al. 2023). Given the absence of defects within the murine retinal vasculature, we returned to the adult murine brain for further analysis of vessel morphometrics.

### KRAS^G12D^ expression in the endothelium increases vessel diameter, volume, and tortuosity in the murine brain

To move the field from a qualitative analysis of the impact of KRAS gain of function on the cerebrovasculature we further interrogated our micro-CT data to establish more quantitative benchmarks on multiple vessel parameters across the entire brain: a task that would be all but impossible using data gathered from 2D imaging modalities. Reconstruction of the micro-CT data using NRECON and 3D volume rendering using CT-VOX allowed us to view the entire murine brain vasculature in three dimensions (**Fig. 5A**). Subsequent quantitative analysis using the open-source software application VesselVio (Bumgarner and Nelson 2022) revealed that there were overall increases in vessel volume (**Fig. 5B**), average vessel diameter (**Fig. 5C**), vessel surface area (**Fig. 5D**), and tortuosity (**Fig. 5E**). To determine if changes in frequency and vessel volume were evenly distributed across different size vessels, we processed this data with CT-Analyzer (CT-AN) (as VesselVio currently lacks this functionality) (**Fig. 5F, Supp. Fig. 6A**). Similarly, we found that the total vessel volume (**Supp. Fig. 6B**) and total surface area (**Supp. Fig. 6C**) were significantly increased in the *ibEC-Kras^G12D^* mice compared to the controls. Furthermore, *ibEC-Kras^G12D^* brains showed a decrease in the frequency of small diameter vessels (e.g. those with diameters <50 μm) and an increase in the number of medium and large diameter vessels (e.g. between 110 and 310 μm) compared to control animals (**Fig. 5G**). Importantly, this same analysis of the whole brain cerebrovasculature in the *ibEC-Kras^G12C^* model detected similar increases in vessel volume, radius, and tortuosity, although changes in vessel frequency were not significant (**Supp. Fig. 7A-G**). Similar to mice expressing G12D, total vessel volume and total surface area were significantly increased in the *ibEC-Kras^G12C^* mice compared to the control animals (**Supp. Fig. 8A-C**). Despite identifying significant differences in vessel diameter, volume, and tortuosity, we wondered if analyzing the entirety of the cerebrovascular network as a single unit masked more dramatic regional changes, particularly in the area harboring the AVM.

**Figure 5.**
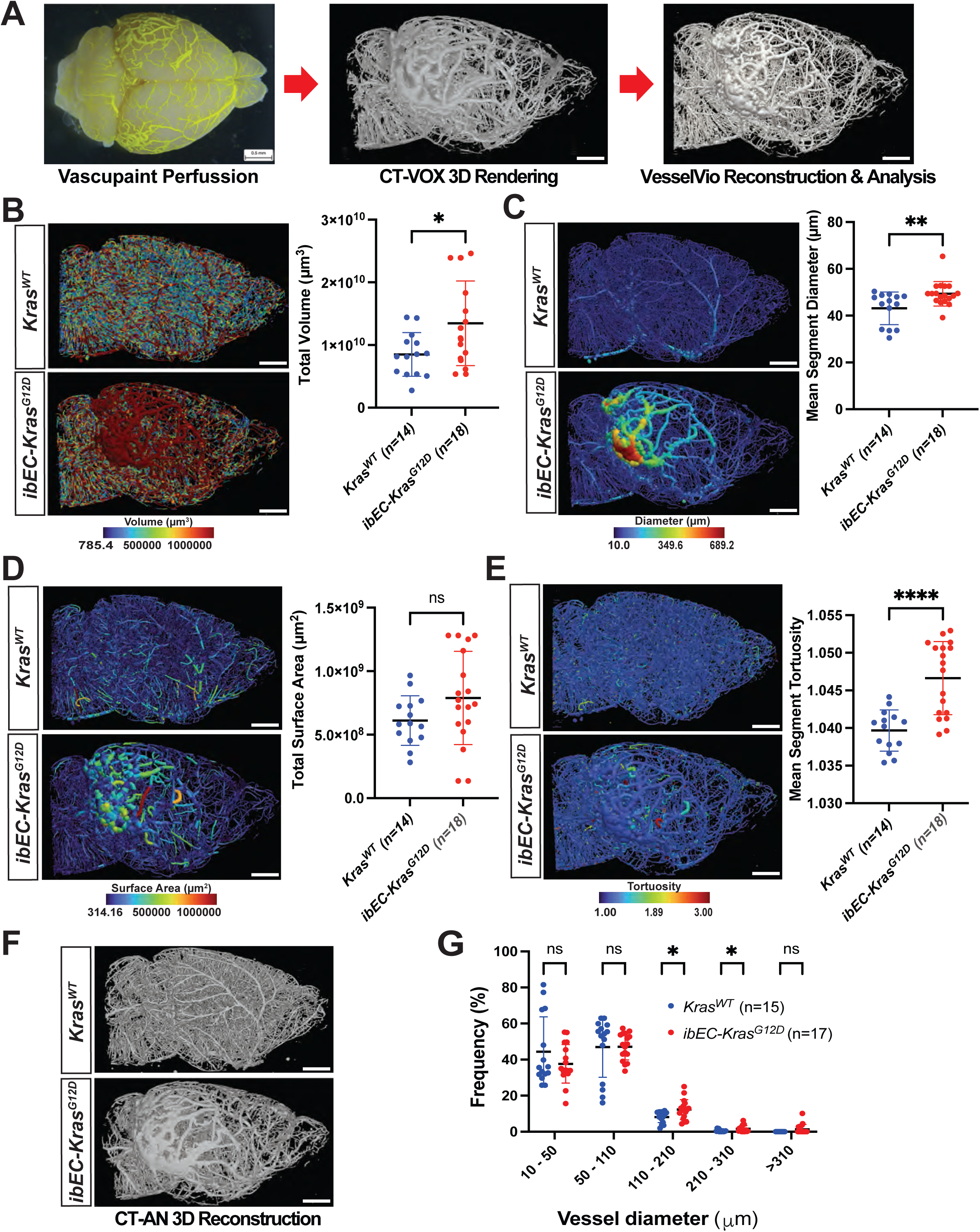
*ibEC-Kras^G12D^* mice feature increased vessel diameter, volume, and tortuosity across the entire cerebrovasculature. (**A**) Pipeline for vessel perfusion and 3D cerebrovascular image reconstruction. The arterial circulation is labelled using Vascupaint contrast agent, followed by 3D micro-CT imaging and 3D rendering using CT-VOX software. Vessel tracing and morphometric analysis are then performed using VesselVio software. (**B**) Sagittal view of representative heat maps from 2 month old adult mouse brains color-coded by vessel volume in (top left) control *Kras^WT^* (n=14) and (bottom left) *ibEC-Kras^G12D^* (n=18) mice. Right side, quantification of total vessel volume for each genotype (each dot represents one unique brain). Scale bar = 2000 μm. *p=0.0312 (student’s T-test). (**C**) Sagittal view of representative heat maps from 2 month old adult mouse brains color coded by vessel diameter in (top left) control *Kras^WT^* (n=14) and (bottom left) *ibEC-Kras^G12D^*(n=18) mice. Right side, quantification of mean segment diameter for each genotype (each dot represents one unique brain). Scale bar = 2000 μm. *p=0.0072 (student’s T-test). (**D**) Sagittal view of representative heat maps from 2 month old adult mouse brains color coded by vessel surface area in control *Kras^WT^* (n=14) and *ibEC-Kras^G12D^* (n=18) mice. Right side, quantification of total vessel surface area for each genotype (each dot represents one unique brain). Scale bar = 2000 μm. p=0.1119 (student’s T-test). (**E)** Sagittal view of representative heat maps from 2 month old adult mouse brains color coded by mean segment tortuosity in (top left) control *Kras^WT^* (n=14) and (bottom left) *ibEC-Kras^G12D^* (n=18) mice. Right side, quantification of mean segment tortuosity for each genotype (each dot represents one unique brain). Scale bar = 2000 μm. ****p<0.001 (student’s T-test). (**F**) Sagittal view of a vessel tracing using CT-AN for 3D reconstruction and analysis following micro-CT imaging. Scale bar = 2000 μm. (**G**) Quantification of the frequency of various diameter vessels in control *Kras^WT^* and *ibEC-Kras^G12D^* mice (each dot indicates one brain from a unique 2-month-old mouse). A student’s T-test was conducted for each comparison. At 10 – 50 μm, p=0.2264; for 50 – 110 μm, p=0.9636; for 110 – 210 μm, *p=0.0169; for 210 – 210 μm, *p=0.0319; for >310 μm, p=0.0672.

Accordingly, the murine micro-CT data was subset into four sections corresponding to the stereotactic coordinates where bAVMs were observed in our earlier analysis (e.g. Bregma +3 to +1, +1 to -1, -1 to +1, and +1 to -5) (**Supp. Fig 9A**). Interestingly, we observed a significant decrease in small diameter vessels (10-110 micron) in *ibEC-Kras^G12D^* mice across the entire rostral to caudal axis of the brain (**Supp. Fig. 9B-E**). Conversely, medium diameter vessels (110-210 micron) were either significantly increased (+3 to +1, -3 to -5) or trended toward an increase (+1 to -1, -1 to - 3). Notably, vessels with diameters ranging between 210-310 microns and larger than 310 microns were evident in the poster region of mutants, but not control animals, although this difference was not significant.

When analyzing these regions <u>o</u>f interest [ROI], we found that many software programs failed to accurately reconstruct vessels within and surrounding the vascular anomaly compared to the initial rendering in CT-VOX, often complicating, or precluding meaningful quantification (**Fig. 6A**). We determined that a recent commercial tool for vessel reconstruction and quantification, Vesselucida, was superior to other programs (i.e. CT-AN and VesselVio) (Bumgarner and Nelson 2022)for reconstructing and tracing vessel networks in the AVM region (**Fig. 6A**).

**Figure 6.**
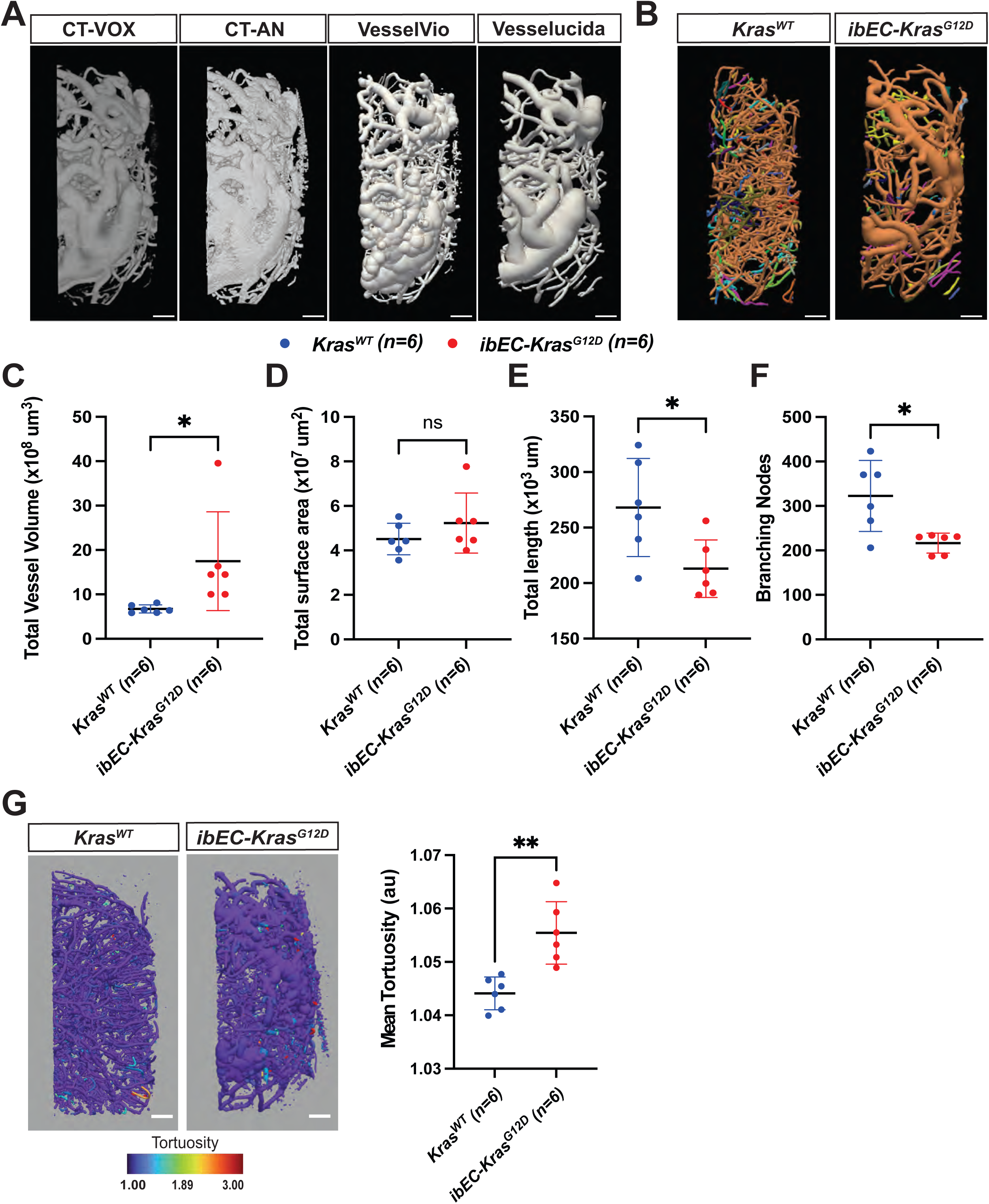
*ibEC-Kras^G12D^*–induced BAVMs feature extensively remodeled vessels. (**A**) Isolation of a region of interest [ROI] containing a bAVM, as opposed to whole brain reconstructions and analysis from the entire mouse brain as seen in Figure 5, after micro-CT imaging and NRECON reconstruction followed by 3D rendering using CT-VOX, CT-AN, VesselVio, and Vesselucida. (**B**) Vessel tracing of the ROI that usually contains a bAVM (Bregma -3 to -5) in *ibEC-Kras^G12D^* mutants. Vessels are color coded to denote directly-connected vessel networks. Scale bar = 500 μm. (**C**) Quantification of vessel volume within the ROI in control *Kras^WT^* and *ibEC-Kras^G12D^* mice (each dot indicates an ROI from a single animal). *p= 0.0398 (Student’s T-test). (**D**) Quantification of total surface area for all vessels within the ROI from control *Kras^WT^* and *ibEC-Kras^G12D^* mice (each dot indicates an ROI from a single animal). p= 0.2753 (student’s T-test). (**E**) Quantification of total vessel length in the ROI from control *Kras^WT^* and *ibEC-Kras^G12D^* mice (each dot indicates an ROI from a single animal). p= 0.0251 (student’s T-test). (**F**) Quantification of branching nodes within the ROI from control *Kras^WT^* and *ibEC-Kras^G12D^* mice (each dot indicates an ROI from a single animal). *p= 0.0106 (student’s T-test). (**G**) Visual heat map reconstruction from VesselVio of vessel tortuosity within the ROI from *Kras^WT^* and *ibEC-Kras^G12D^* mice on the left and quantification on the right (each dot indicates an ROI from a single animal). Scale bar = 500 μm. **p= 0.0018 (Student’s T-test).

Using this software, we found that total vessel volume was significantly increased in the [ROI] that most frequently contained the bAVM in mutants compared to the same region in control animals (**Fig. 6B, C**). While the total surface area of the vessels trended toward an increase in mutants, this difference was not significant between the two genotypes (**Fig. 6D**). However, total vessel length in the region of interest was significantly lower in mutants compared to controls (**Fig. 6E**). In addition, the number of branching nodes, an indicator of vessel density, was significantly lower in the bAVM area in mutants (**Fig. 6F**). Using Vesselvio, we quantified vessel tortuosity in the ROI (as Vesselucida currently lacks this functionality). Notably, vessel tortuosity in the Bregma -3 to -5 area was significantly increased in *ibEC-Kras^G12D^* mice compared to their control littermates (**Fig. 6G, H**). Similar to KRAS^G12D^, endothelial-specific expression of KRAS^G12C^ significantly increased vessel volume and tortuosity within the bAVM ROI (**Supp. Fig. 10A-G**).

### Aging does not potentiate active KRAS-induced morphological changes in the murine cerebrovasculature, but increases mortality

Next, we wanted to determine if aging impacted KRAS-driven changes in the cerebrovasculature. Following tamoxifen injection at P1, both *Kras^WT^* and *ibEC-Kras^G12D^* mice were aged to either 7 or 14 months of age. We previously reported that survival of *ibEC-Kras^G12D^* mutants and littermate controls was indistinguishable through the first two months of life(Fish et al. 2020). However, by 12 months of age more than 50% of *ibEC-Kras^G12D^* mice (n=27) had died compared to just 25% of controls (n=20) (*p= 0.0186), with *ibEC-Kras^G12D^* yielding a hazard ratio of 0.3064 (Mantel-Haenszel) (**Supp. Fig. 11A**). Of note, bAVM incidence was greater at 2 months (n=15/22, 68.18%) than at 7 months (n=6/13, 46.15%) (**Supp. Fig. 11B,C**). bAVM incidence decreased at 14 months (n=2/6) to 33.33% (**Supp. Fig. 11D,E**), when the probability of survival was near 25%. We postulate that these differences in bAVM incidence may be due to reduced survival of animals with severe bAVMs as they age, although the inability to perfuse the vasculature post-mortem precluded testing this hypothesis. Alternatively, this difference could be due to a sampling error given the small number of mice present at this stage.

Next, we used micro-CT imaging and analysis to determine how aging impacted KRAS-induced changes in vessel structure and remodeling (**Supp. Fig. 12A**). Morphometric analysis of the ROI containing the bAVM showed a significant increase in the number of branching nodes, a proxy for vessel density, in the mutants compared to the controls at 7 months of age (**Supp. Fig. 12B,C**). Similarly, total vessel length, surface area, volume, and vessel tortuosity were all significantly increased in the ROI of mutants harboring a bAVM compared to the same region in control animals (**Supp. Fig. 12D-G**).

To address whether vessels underwent pathologic remodeling with aging, we quantified the percent change in vessel length between mutant and wild-type littermates at 2 and 7 months. A similar analysis was performed to determine the percent change in surface area, total vessel volume, and tortuosity. Except for vessel length, significant differences were not apparent when comparing changes in vessel morphometry at 2 and 7 months (**Supp. Fig 12H-K**).

### *ibEC-Kras^G12D^* mutant mice feature altered behavior, cognition, and motor function

While most sporadic bAVMs tend to be clinically asymptomatic(Derdeyn et al. 2017), some patients develop seizures, headaches, visual disturbances, ataxia, aphasia, and muscle weakness(Lantz and Meyers 2008). A previous study showed that adult mice transduced with AAV over-expressing *Kras*^G12V^ in ECs (and potentially other neuronal cell types) which feature multiple bAVMs at a high frequency/penetrance (100%), present with sensory, cognitive, and memory deficits, as well as motor behavior dysfunction(Park et al. 2021). Given that this virus readily transduced non-endothelial cells in the brain (e.g. neurons) (**Supp. Fig. 1**), whether the reported neurobehavioral defects were due to mutant KRAS activity within, or outside, of the endothelium in this AAV model remains unclear.

To define the impact of endothelial gain of KRAS function on murine cognition and behavior, we first examined long-term and spatial memory using the <u>n</u>ovel <u>o</u>bject recognition (NOR) test(Lueptow 2017). We observed a significant difference in the NOR index in bAVM-bearing *ibEC-Kras^G12D^* compared to their control littermates(**Supp. Fig. 13A**). Similarly, there were significant differences in the discrimination index at 7 months of age in mutants with bAVMs compared to controls (**Supp Fig. 13B**), suggesting that endothelial *Kras^G12D^* gain of function induced bAVMs impaired working memory at 7 months of age

Next, we examined general activity, anxiety, and exploratory behavior in *ibEC-Kras^G12D^* and control littermate mice using the open field activity test(Perusini and Fanselow 2015; Robinson et al. 2019). No significant differences in the duration of time spent in the center of the field at 7 months of age were evident between the two genotypes, suggesting *ibEC-Kras^G12D^* mice do not experience greater anxiety than control littermates (**Supp. Fig. 13C**).

Finally, we examined fine motor coordination using the parallel rod footfall test(Kamens and Crabbe 2007). Notably, mutants exhibited significantly more foot falls than control littermates at 7 months of age, also regardless of bAVM status, suggesting endothelial-specific expression of mutant KRAS induced bAVMs impaired fine motor coordination (**Supp. Fig. 13D**). Gross abnormalities in gait are associated with conditions affecting motor function in Parkinson’s disease, spinal cord injury and stroke(Timotius et al. 2023). Quantification of stride length via catwalk analysis (Timotius et al. 2023; Zheng et al. 2023) showed that gait is indistinguishable between the two genotypes at seven months of age, suggesting that no overt abnormalities in gait or gross motor coordination (e.g. ataxia) were evident in *ibEC-Kras^G12D^* animals (**Supp. Fig. 13E**). Further analysis of natural step length, stride duration, paw pressure distribution, and other parameters revealed that the duration of the walk, the number of steps, average speed, and cadence were not significantly different in *ibEC-Kras^G12D^* compared to control littermates (**Supp. Fig 14A-E**). Collectively, these results suggest KRAS gain-of-function in the endothelium and bAVM impairs learning and memory, as well as fine motor coordination, without impacting natural locomotion.

### Inhibition of MEK activity normalizes KRAS-induced bAVMs

Recent work showed that initiating MEK inhibitor (MEKi) treatment concurrently with AAV-transduction of mutant *KRAS^G12V^* – in effect pre-treating before endothelial cells begin expressing mutant KRAS – prevents bAVM induction, while treating early postnatal pups – a period when AVMs are not detectible (Fish et al. 2020) – normalizes vascular dysplasia in mice(Park et al. 2021; Nguyen et al. 2023). However, the clinical relevance of these studies is difficult to interpret, as most patients with sporadic bAVM are diagnosed at 25 years of age. Thus, treatment prior to lesion formation (i.e. before a diagnosis of bAVM is made) is not possible. Instead, the goal of pharmacological intervention would be to stabilize or regress established lesions after their identification. Accordingly, we initiated treatment at 1 month of age in our sporadic mouse bAVM model to test if MEKi treatment could disrupt lesion progression or reverse these anomalies. Mirdametinib (PD0325901) was chosen due to its excellent IC50, and its ability to traverse the blood brain barrier and block ERK phosphorylation across multiple regions of the brain in vivo (unlike Mekinist/Trametinib, SL327 or U0126)(Brown et al. 2007; Barrett et al. 2008; Papale et al. 2016).

We injected *ibEC-Kras^G12D^* pups with tamoxifen at P1, then began treatment 30 days later with either vehicle (n = 22) or mirdametinib (PD0325901) (n = 20) (20 mg/kg p.o., daily) for 30 days (**Fig. 7A**). Mirdametinib did not affect the animal’s weight over the duration of the treatment window, compared to vehicle (**Supp. Fig. 15**). Representative phase microscopy images revealed a normalized vasculature in 2-month-old *ibEC-Kras^G12D^* MEKi treated mice (n=20), with decreased incidence and number of bAVMs per mouse, and fewer of the large, dilated shunts characteristic of vehicle-treated *ibEC-Kras^G12D^* mutant mice (n=22) (**Fig. 7B-D**).

**Figure 7:**
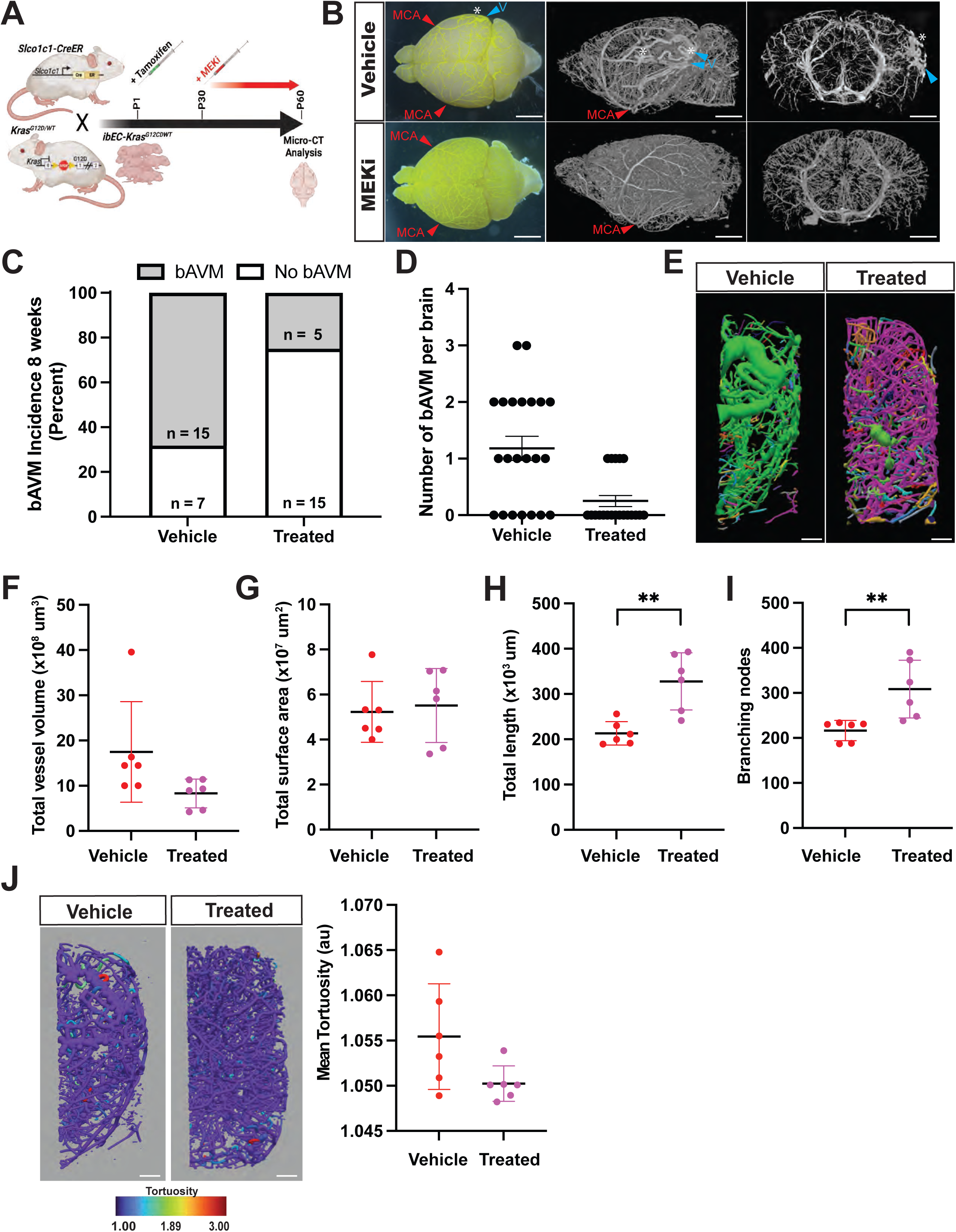
MEK inhibition prevents bAVMs in *ibEC-Kras^G12D^* mice. (**A**) Experimental design. *Slco1c1-CreER;Kras^G12D/+^* pups were given tamoxifen at P1, then at 1 month of age treated daily with either vehicle or MEKi (Mirdametinib/PD0325901) for 1 month. At 2 months of age, animals were perfused with Vascupaint^TM^ contrast agent and imaged by micro-CT. (**B**) Representative phase microscopy (far left column, dorsal view, scale bar = 500 μm) and micro-CT (middle column, sagittal view, scale bar = 500 μm) images of the mouse brains at 2 months of age from following Vascupaint^TM^ perfusion. Red carets and MCA denote the mid cerebral artery. White asterisk denote arteriovenous shunts (bAVMs). Blue carets denote the presence of dilated and tortuous draining veins. Far right column shows a 2,150 μm thick virtual “slab” section and reconstruction of the ROI that typically contains the bAVM (Bregma -3 - -5), coronal view, from the same brains. Note the cortical localization of the arteriovenous malformation in the vehicle treated animal and the absence of bAVM in the MEK inhibitor treated animal. (**C**) Quantification of bAVM incidence at 2 months of age in *ibEC-Kras^G12D^* mice after treatment with vehicle control or MEKi for 1 month. (**D**) Quantification of the total number of bAVMs per *ibEC-Kras^G12D^* mouse brain at 2 months of age. (**E**) Vesselucida reconstruction of the ROI, -3 to -5 Bregma, where bAVMs are most frequently detected in *ibEC-Kras^G12D^* mice. Vessels are color coded to denote directly-connected networks. Scale bar = 500 μm. (**F**) Total vessel volume within the ROI is decreased in *ibEC-Kras^G12D^* treated with MEKi (n=6) compared to vehicle control treated animals (n=6), but it is not significant (p=0.0804, student’s T-test). (**G**) Total surface area within the ROI does not differ between *ibEC-Kras^G12D^* mice treated with MEKi (n=6) or vehicle control (n=6) (p=0.7523, student’s T-test). (**H**) Total vessel length within the ROI is significantly increased in *ibEC-Kras^G12D^* mice treated with MEKi (n=6) compared to vehicle control (n=6) animals (*p=0.0021, student’s T-test). (**I**) The number of branching nodes is increased in MEKi treated animals (n=6) compared to vehicle control (n=6) *ibEC-Kras^G12D^* mice (*p=0.0077, Student’s T-test). (**J**) VesselVio reconstruction of the ROI containing an bAVM. Vessels are color coded to denote directly-connected vessel networks. MEKi treated mice show decreased tortuosity (p=0.0654, student’s T-test) compared to control treated animals.

Morphometric analysis of micro-CT data from the posterior region of the brain where bAVMs are typically located showed that the decrease in the frequency of small diameter vessels (10 – 110 μm) evident in *ibEC-Kras^G12D^* mice compared to control animals was partially rescued with MEKi treatment, as was the increase in medium size vessels (110 – 210 μm) (**Supp. Fig 16A, B**). Morphometric analysis of the ROI showed that MEK inhibition either ameliorated, or trended towards normalizing, multiple other differences in vessel remodeling, including overall vessel volume, vessel length, the number of branching nodes, and tortuosity (**Fig. 7F-J**). In *Kras^WT^* control littermates, MEKi treatment had no significant effect compared to vehicle treatment on vessel volume, surface area, length, branching, and tortuosity (**Supp. Fig. 17**). Collectively, these data show that MEKi normalized pathologic vessel remodeling and reduced bAVM incidence in *ibEC-Kras^G12D^* mice.

### Pharmacological Blockade of MEK1 Activity May Prevent bAVM Progression in Pediatric Patients

Widespread ERK activation may be a unifying feature of sporadic bAVM, regardless of the variant allele frequency or underlying somatic mutation(Nikolaev et al. 2018). This, combined with the fact that more than half of all adult bAVMs feature activating mutations in *KRAS(Nikolaev et al. 2018)*, and our previous animal studies showing that MEK inhibition can reverse shunts in embryonic zebrafish (Fish et al. 2020) and data presented herein, suggest that MEK may be a rational therapeutic target in bAVM patients, particularly in those with large AVMs not amenable to surgical intervention or radiation therapy. This particular patient population has no treatment options available, highlighting the importance of identifying novel treatment modalities to alter their clinical course.

Thus, in our clinical practice, we have applied for compassionate use of the MEK inhibitor, trametinib, for severe pediatric patients at Texas Children’s Hospital. Pediatric patients with brain AVMs deemed either too large for radiation therapy or untreatable with surgical resection (due to large size, eloquent location, or usually both) were treated with trametinib. Three pediatric cases (**Fig. 8**) have shown stable disease with little to no progression since beginning treatment, although this follow-up timeframe is truncated and further follow-up is needed. As this treatment was under compassionate use, and the optimal therapeutic dosing is unknown, the starting dose was low with vigilant monitoring for side effects, including drug-induced dermatitis, abdominal discomfort, rhabdomyolysis, and other issues.

**Figure 8:**
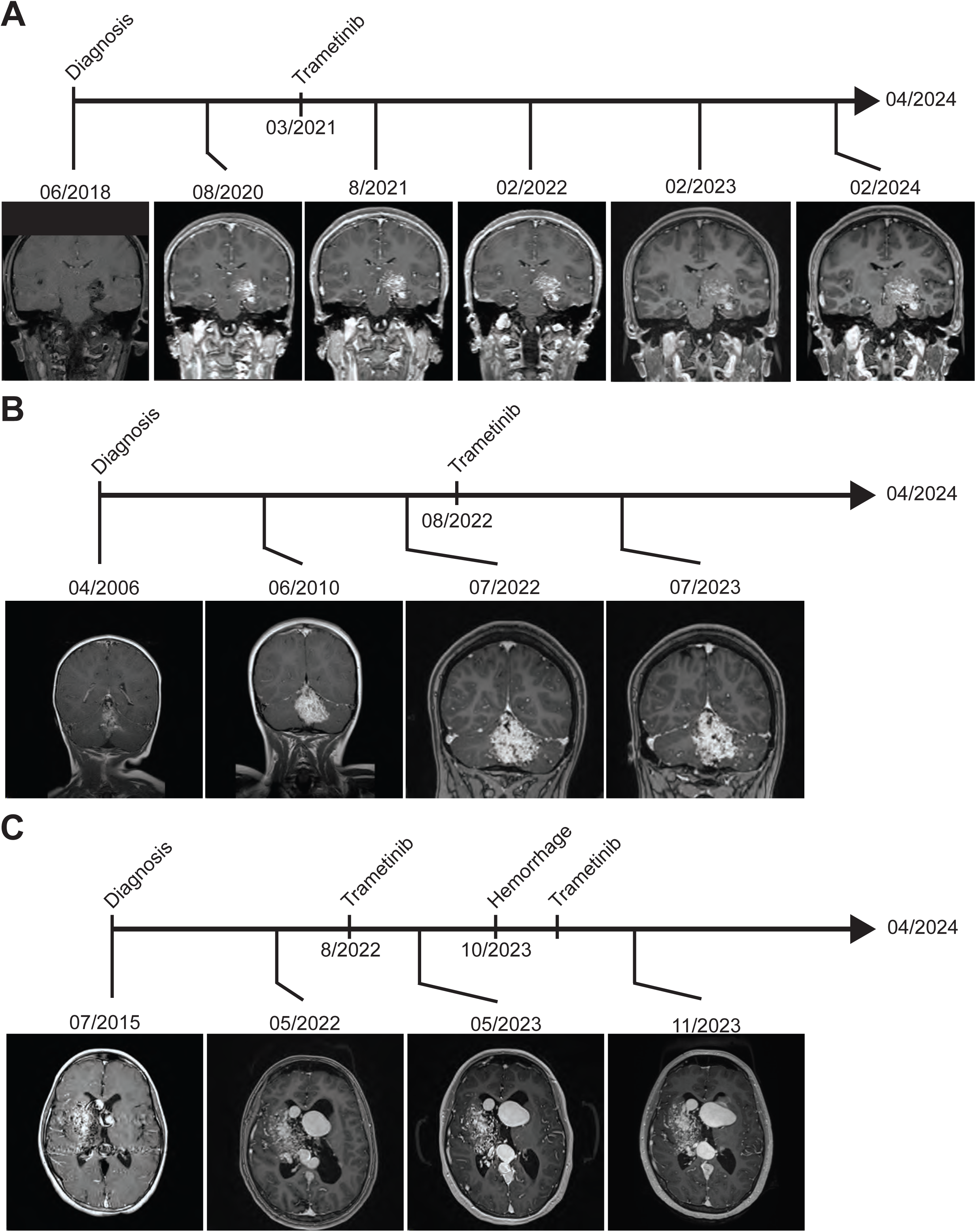
MEK inhibition may stabilize bAVMs in pediatric patients. (**A**) Patient A, a now 14 year old female, presented with a left thalmic AVM in 2018 as determined by MR imaging. She was briefly lost to follow up for several years, but repeat imaging in 2020 revealed progression of the AVM, now a Spetzler-Martin Grade 4. Trametinib treatment (0.5 mg p.o., daily, increased after 3 months to 1 mg p.o. daily) was begun 3/25/2021. Follow up MRI imaging in February 2022 through February of 2024 showed relative stability. (**B**) Patient B, a now 19 year old female, who possessed a vermian cerebellar AVM originally diagnosed in 2006, showed lesion progression and enlargement of the nidus in 2010. Their lesion was consistent with a Spetzler-Martin Grade 4 AVM in July of 2022 and trametinib treatment was initiated in August of 2022 (0.5 mg p.o., daily, increased after 3 months to 1 mg p.o. daily), with relative stability noted in a follow up scan in July of 2023. (**C**) Patient 3 a now 19 year old female, presented with a large right basal ganglia and thalmic AVM in 2015, but was lost to follow up until 2022 when MR imaging was repeated and revealed substantial progression in the AVM nidus and enlargement of deep draining veins, now a Spetzler-Martin Grade 5 AVM. Trametinib was initiated in August of 2022 (1 mg p.o. daily) and follow up in May of 2023 showed lesion stabilization and no significant progression. In Oct 2023, she unfortunately had a hemorrhage, but recovered post-bleed and was restarted on trametinib (1 mg p.o. daily), with recent imaging in November 2023 showing continued relative stability of her AVM.

Patient A, an 8-year-old female at initial presentation (now 14 years of age), presented with a left thalamic AVM, initially diagnosed in 2017. Follow-up imaging by MRI in 2020 showed progression of the AVM with nidal enlargement and trametinib was initiated (0.5 mg p.o., daily, increased after 3 months to 1 mg p.o. daily). Since beginning treatment in March of 2021, subsequent imaging in February of 2022 and February of 2024 has shown relative stability in the lesion. Patient B, a now 19 year old female, who had a vermian cerebellar AVM, originally diagnosed in 2006 at age 2, was closely followed for several years with serial imaging and no interventional treatment. Lesion progression and enlargement of the nidus was noted in 2010, and trametinib treatment (as above) was begun in August of 2022, with relative stability in imaging parameters since that time. Patient C, a now 19 year old female, who presented initially at 11 years of age with a large right basal ganglia and thalamic AVM in 2015, but was subsequently lost to follow up for several years. Repeat imaging in 2022 showed substantial progression in the nidus and enlargement of deep draining veins. Trametinib was initiated in July 2022, and follow up one year later showed lesion stabilization and no significant progression. However, in October 2023, Patient 3 had a hemorrhagic stroke, but was still deemed ineligible for surgical or endovascular intervention and trametinib was continued, but with an increased dose of 1 mg p.o. per day.

While the *KRAS* variant status of each of the lesions in these patients is unknown, these initial case study results are encouraging, although they are anecdotal at this point given the limited longitudinal follow up since initiating treatment. However, given the current availability of three FDA-approved MEK inhibitors with relatively strong safety profiles and demonstrated efficacy in treating solid tumors(Han et al. 2021; Bouffet et al. 2023), follow up clinical trials for novel drug indications of these agents in bAVM is hopefully on the horizon.

## DISCUSSION

Since the discovery in 2018 that more than 50% of sporadic bAVM cases in humans contain gain-of-function genetic variants within the *KRAS* gene(Nikolaev et al. 2018), and the follow-up demonstration that constitutively active Kras^G12D^ is sufficient to induce bAVMs in mice, while Kras^G12V^ or Kras^G12D^ variants were sufficient to induce AVMs in zebrafish, our understanding of the mechanisms driving these potentially devastating vascular anomalies has undergone a renaissance. The initial human genetic results have since been validated by other groups(Goss et al. 2019; Hong et al. 2019; Oka et al. 2019; Priemer et al. 2019; Gao et al. 2022), while another report showed that AAV-mediated transduction of human KRAS^G12V^ was sufficient to induce bAVMs in adult mice(Park et al. 2021).

In the current study, we confirm that endothelial-specific expression of KRAS^G12D^ robustly induces bAVM in mice, and extend these observations to define the phenotypic consequences on the murine cerebrovasculature in the setting of KRAS gain-of-function. Micro-CT imaging and analysis of 3D imaging data of the entire brain revealed that mutant KRAS^G12D^ yielded bAVMs predominantly in the caudal (i.e. posterior) portion of the brain, between Bregma -3 to -5, with an average of 1 lesion per mouse (up to 3 lesions per animal), and that these vascular anomalies were localized primarily to the cortex. Morphometric analysis of the entire cerebrovasculature revealed that KRAS expression led to increased vessel volume, radius, and surface area, and that vessels in the brain overall were more tortuous. Detailed analysis of the region that usually contains the bAVM showed that in addition to the previous changes, vessel density was decreased, and fewer small diameter vessels were present at 2 months of age. These changes persisted in aged mice, with the exceptions that unlike at 2 months of age, vascular density and vessel length were increased in aged animals.

While it is unknown whether KRAS^Q61H^, another variant identified in bAVM patients(Nikolaev et al. 2018), is sufficient to induce lesion formation, using both zebrafish and mice we have shown that two mutant variants affecting Glycine 12 of KRAS (KRAS^G12D^, KRAS^G12V^) are sufficient to induce bAVM formation in vivo(Fish et al. 2020). Multiple reports show that *KRAS^G12D^* is the most common variant in sporadic bAVM, and interestingly Gao and colleagues suggested that *KRAS^G12V^* (∼2%) has a greater allelic burden than *KRAS^G12D^* (∼1%)(Gao et al. 2022), raising the question of whether KRAS^G12D^ is a more pathogenic variant in bAVM than KRAS^G12V(Gao et al. 2022)^. Notably, in vitro studies in epithelial cancer cells suggest different KRAS variants exert unique effects on proliferation, selection, and downstream signaling(Monticone et al. 2008). However, a clear association between mutation allelic burden and bAVM severity phenotype (measured by age at resection, lesion size, or age at first hemorrhage) has not been established in humans or animals. Provocatively, while our initial report identified G12D, G12V and Q61H variants in bAVM(Nikolaev et al. 2018), we failed to identity other activating KRAS mutations in our patient cohorts. However, a subsequent study identified G12C variants in bAVM(Priemer et al. 2019). Both G12D and G12C variants are of particular interest given the rapid progress in identifying potential inhibitors targeting these two isoforms in the field of oncology(Canon et al. 2019; Wang et al. 2022); however, their clinical utility remains to be determined(Blaquier et al. 2021; Liu et al. 2022). Nonetheless, we were curious whether the G12C mutant variant is sufficient to drive bAVMs in mice, and whether these mice would feature defects similar to those of Kras^G12D^ mice, as this would suggest G12C mutations should also be given equal scrutiny for targeted diagnostic sequencing in patients.

We found that KRAS^G12C^ led to approximately 2 lesions per animal, with a somewhat wider spatial distribution compared to KRAS^G12D^ gain-of-function, with bAVMs occurring in the rostral/anterior (Bregma +3 to +1) but predominantly in the caudal/posterior (-3 to -5) regions of the adult mouse brain. Of note, the posterior location was where most lesions also occurred in the KRAS^G12D^ mice. Similar to KRAS^G12D^, anomalies were most frequently detected in the cerebral cortical region.

The localization of lesions in our KRAS^G12D^ and KRAS^G12C^ models contrast those where mice were transduced with *AAV2-BR1-CAG-hKRAS^G12V^-WPRE(Park et al. 2021)*. Although the precise location within the brain was not reported(Park et al. 2021), approximately 91% of these mice developed a bAVM in the olfactory bulbs. The virtual absence of olfactory bulb involvement in the G12D or G12C genetic models may be due to the synthetic/artificial CAG-promoter driven expression in the AAV system which likely results in non-physiologic levels of mutant *KRAS* transcripts, and viral transduction outside of the brain endothelium, versus the endothelial-restricted activity of the *Slco1c1-CreER* driver and *Kras^G12D^* and *Kras^G12C^* transcripts being regulated by the endogenous *Kras* promoter in these genetic models. RNA-scope revealed that *Kras* transcripts appear to be enriched in the cortical endothelium compared to the olfactory bulb. Differential Cre recombinase activity in the olfactory bulb compared to the cortex in the genetic models also does not explain these differences, as *Slco1c1-CreER* activity in the adult mouse brain is uniform across the rostral to caudal axis following tamoxifen administration at P1(Fish et al. 2020). Other variables, such as the age of the mice at viral transduction (5 weeks of age) vs tamoxifen administration (P1) likely do not explain this difference either, as Cre-mediated recombination of the *Kras* locus would persist into the adult CNS endothelium. Whether regional heterogeneity, specifically compartmental differences in gene expression and cell signaling cascades (Dubrac et al. 2023) explains the differential frequency of bAVMs across unique regions of the brain warrants additional, further study.

In humans, we observed that bAVMs are located primarily in the lobes of the brain which are the subdivisions of the cerebral cortex. Specifically, the frontal lobe was the most common site for bAVM, followed by the midline structures and the temporal lobe, with approximately 25% of these vascular anomalies occurring in the parietal-temporal lobe in human patients. Notably, the *KRAS* mutational status for the patients used for our retrospective case study are unknown, but future studies involving endovascular biopsy followed by genetic testing could determine whether *KRAS*-induced lesions exhibit any specific anatomical preference or susceptibility to intracranial hemorrhage.

For the first time, we have quantified the morphological consequence of KRAS gain of function in the CNS vasculature in both the *ibEC-Kras^G12D^* and *ibEC-Kras^G12C^* models using a novel imaging modality, namely micro-CT. Our 3D analysis of the entire brain vasculature determined that the frequency of small diameter vessels decreases while the number of large diameter vessels increases in mutant KRAS animals. Notably, these changes are even more drastic within the region that usually contains the bAVM. Furthermore, there was an increase in vessel tortuosity in the bAVM region as well. Although overall survival was significantly diminished in KRAS^G12D^ mutant mice, time did not appear to exacerbate the vascular morphological differences, as changes in vessel volume, vessel surface and tortuosity in the vessels at 7 months of age were comparable to the those observed at 2 months. This study represents the first side-by-side whole-brain and bAVM-specific morphometric analysis of vessel morphology in a murine bAVM model. These new benchmarks will create a standard for future comparative studies with other bAVM mouse models, allowing others to identify similarities, as well as distinguishing features, of various somatic and germline murine models that feature bAVMs.

Critically, bAVM patients can feature pronounced neurocognitive deficits(Mahalick et al. 1991; Choi et al. 2009). This may be due in part to increased blood flow through the shunt and the resultant diminished circulatory flow through adjacent capillary vessels. This hypoperfusion may induce focal atrophy and cell death in the parenchymal area adjacent to the anomaly(Wang et al. 2021), although evidence for this so called “steal” concept is contentious(Okabe et al. 1983; Spetzler and Zabramski 1988; Mast et al. 1995; Meyer et al. 1998; Meyer et al. 1999). In a blinded, cross-sectional study, 71.4% of bAVM patients (n=70) exhibited neurocognitive deficits, without any detectable neurological features(de Souza Coelho et al. 2019). This same report noted a significant association between bAVMs in the temporal lobe and memory deficits, in agreement with an established role for the temporal lobe in regulating memory and language processing. Notably, in this study, Spetzler-Martin (SM) grade did not correlate with neurocognitive deficits in patients with unruptured bAVMs. However, SM grade was directly correlated with deficits in working memory in patients with ruptured bAVMs(de Souza Coelho et al. 2019).

In humans, the posterior parietal cortex (PPC) mediates sensory functions, such as spatial attention, spatial awareness, and sensory integration; but it is also involved in movement and decision making(Andersen and Cui 2009). This is similar to mice, in which the PPC is involved in sensory and multisensory processing, navigation, motion planning, and decision-making(Lyamzin and Benucci 2019). In humans, bAVMs located in the parietal cortex have been associated with deficits in sensory integration(Deng et al. 2020), however, to our knowledge larger retrospective studies focused on correlating anatomical location and AVM grade with sensory function have not been undertaken. Similarly, whether sporadic bAVM correlates with decreased neurocognitive function in animal models is unclear.

*AAV-KRAS^G12V^* mice have impaired olfactory function, cognitive and memory deficits, as well as loss of grip strength(Park et al. 2021). That bAVMs located within the olfactory bulb lead to neurocognitive and motor deficits is somewhat surprising, since learning and memory, as well as coordination, are typically controlled by the hippocampus and somatosensory/motor cortex, although the high reliance on olfaction in mice could conceivably impact other neurocognitive behaviors. As our *Kras^G12D^* and *Kras^G12C^* models rarely featured bAVMs in the olfactory bulb, we instead focused on neurologic functions that could be impacted by altered circulation in regions where bAVMs were observed (e.g. the cortex, hippocampus, or striatum). Endothelial-specific recombination of *Kras^G12D^* hindered spatial learning and memory function at 7 months of age and attenuated fine motor function. Importantly, these defects were evident in mice regardless of whether they contained an obvious bAVM, suggesting that endothelial-specific expression of mutant KRAS negatively impacts neurovascular health.

In cancer, therapies for *KRAS*-mutant tumors frequently target downstream signaling molecules, such as MEK, due to the historical difficulties in targeting RAS(Jessen et al. 2013; Merchant et al. 2017). While the recent identification of variant specific KRAS inhibitors may signal a breakthrough in this area, the efficacy of these in the clinic remains unclear(Ricciardelli et al. 2023). Previously, we demonstrated that MEK inhibition both prevents, and regresses, KRAS-driven AVMs in an embryonic zebrafish model(Fish et al. 2020). Using a model that we previously developed, where *Kras^G12D^* is recombined throughout the postnatal endothelium (e.g. *Cdh5-PAC-CreER*; *Kras^G12D/+^*), Nguyen and colleagues showed that *Kras^G12D^* induced vascular dysplasia in the postnatal intestine, heart, liver, and lung(Nguyen et al. 2023). They went on to show that treatment with the MEK inhibitor Trametinib alleviated these defects in postnatal mice. However, as we initially reported, this genetic model does not exhibit bAVMs prior to extensive postnatal demise by P21. Thus, whether Trametinib prevents, or regresses, bAVM formation in a genetic KRAS gain-of-function model of sporadic bAVM remained unknown. Additionally, Trametinib treatment in adult mice beginning 1 day after viral transduction with *AAV-BR1-hKRAS^G12V^* led to a reduction in bAVMs incidence when they were examined 6 weeks later(Park et al. 2021), but again, this was done in the context of prevention, rather than treatment of existing lesions.

To extend the translational impact of these studies, we wanted to instead determine if MEK inhibition could normalize KRAS^G12D^-induced pathological remodeling of the cerebrovasculature, or potentially regress sporadic bAVMs. Thus, we turned to our genetic mouse model. Strikingly, we show that Mirdametinib treatment one month after induction of mutant KRAS expression reduced bAVM incidence by more than 50% in *ibEC-Kras^G12D^* mice, and normalized multiple morphometric changes in the cerebrovasculature. We extended these mouse studies to the clinic, showing that Trametinib, an FDA-approved MEK inhibitor, may be effective at slowing lesion progression in pediatric patients. Future clinical trials are expected to determine whether MEK inhibitors may be of use in treating these patients.

Taken together, we demonstrated that two KRAS mutations, G12D and G12C, are sufficient to induce bAVMs in mice. We further characterized the morphogenetic consequences of KRAS gain of function in the murine cerebrovasculature and compare these features (such as frequency and location) to what is seen in human patients. Then, we determined MEK activity is required for KRAS-driven bAVM formation in mice. Furthermore, we provide a pilot cohort of pediatric human patients that suggests MEK inhibition may be an effective therapeutic in humans. Our data enriches our understanding of KRAS-driven bAVM formation and suggests MEK inhibition as a potential therapeutic treatment.

## MATERIALS and METHODS

### Human Data

#### Study Approval

All human data was collected at Texas Children’s Hospital and Baylor College of Medicine, with informed consent, under IRB protocol H-51515. All data and patient information were deidentified.

#### Magnetic Resonance Imaging

MR imaging was performed on each patient using a standard full brain MRI with and without contrast and MR angiography of the head with and without contrast. Views shown within the manuscript (e.g. coronal for patient Figure 9 panels A and B, and axial for panel C) were chosen to provide the best visualization of AVM size at maximal diameter, but all sequences were done for all patients. MRI brain T1 with contrast are the sequences shown for all 3 patients.

### Murine Experiments

#### Study Approval

All mouse experiments were approved by the Institutional Animal Care and Use Committee (IACUC) at Baylor College of Medicine (protocol AN-7731), the University of Virginia ACUC (#4446) and the University of Texas Health Sciences Center at Houston (UTHSC) IACUC (#AN-23-0052).

#### Mouse Lines Used

The *lox-stop-lox Kras^G12D^* mutant line, *Kras^G12D/WT^* (aka *Kras^lsl-G12D^*, *Kras^tm4Tyj/WT^*, MGI: 2429948). was rederived at the Genetically Engineered Mouse Core (GEM Core) at Baylor College of Medicine using sperm acquired from Jax labs; and *Slco1c1(BAC)CreER* (aka *Slco1c1-iCre/ERT2^Mrks^*, MGI: 5301361), from Dr. Markus Schwaninger (University of Lübeck), were kindly provided by Dr. Mark Kahn (University of Pennsylvania). For breeding purposes, *Kras^G12D/WT^* mice were bred onto an *R26^RFP^* (specifically *Rosa26^CAG-lsl-TdTomato^, R26^ai14^*, MGI: 3809524) background. Females heterozygous for the conditional mutant *Kras* allele and homozygous for a Cre-dependent fluorescent reporter (i.e. *Kras^lsl-G12D/WT^*; *R26^RFP/RFP^*) were crossed to males harboring *Slco1c1-CreER* that were wildtype for *Kras* (e.g. *Slco1c1-CreER*; *Kras^WT/WT^*) to generate experimental litters (see below for more details). A similar strategy was undertaken for the *Kras^G12C^* allele (*B6;129S4-Kras^em1Ldow/J^*; MGI: 6281558). All mice were genotyped by PCR. Please see **TABLE I** for primers and details.

**TABLE I.**
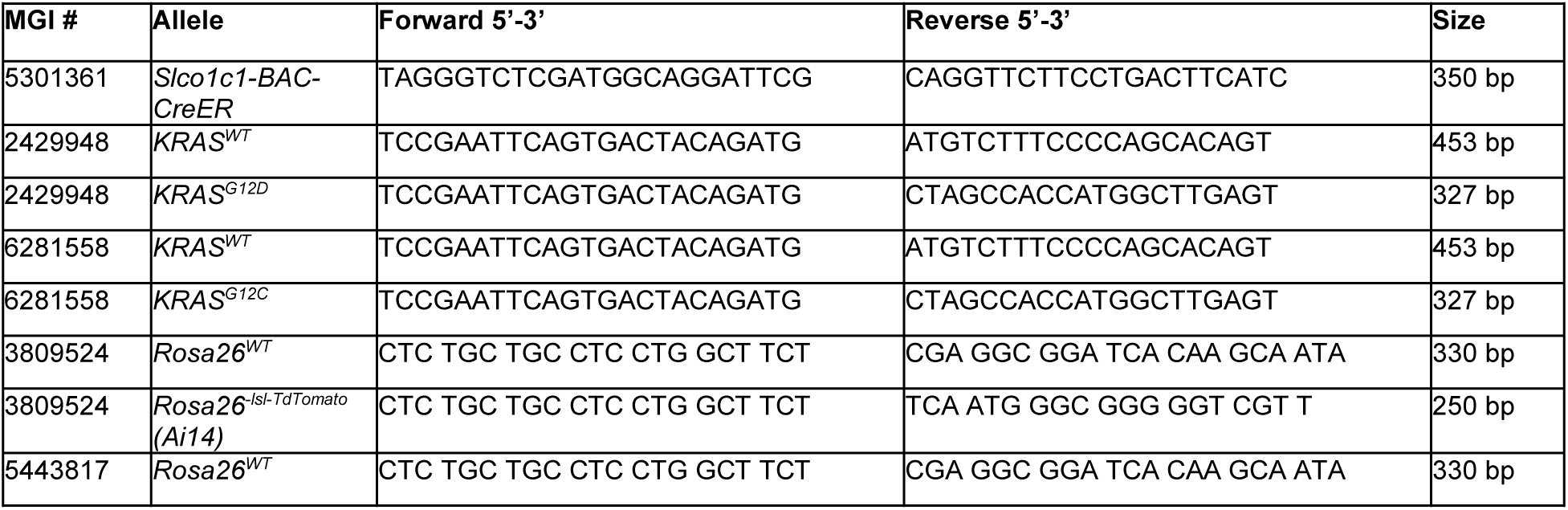
Mouse lines used and gene-specific PCR genotyping primers.

#### Randomization of Animals for Treatment Groups, Experimental Blinding, and Sex as a Biological Variable

For all postnatal, brain endothelial experiments, all pups, regardless of sex or genotype, were injected with tamoxifen at P1. Animals were then marked by toe-clip after injection (and later by a metal ear tag with a unique alphanumeric identifier at P21) and that genetic material was used later for PCR-based genotyping. Images for analysis of any phenotype were recorded by a person blinded to the animal’s genotype. These images were then scored or quantified by another person blinded to the animal’s genotype, and then each animal’s unique identity was then matched to their respective genotypes. For adult, pan-endothelial gain of function studies, all littermates of each genotype, regardless of sex, were injected with tamoxifen and then later imaged or processed by one individual at the time of harvest, while the images were scored by a second person blinded to the animal’s identity. All littermates, both males and females, were used for experimental analysis as described below. We did not assess sex-specific differences in this study due to the small sample size and lack of documented sexual dimorphism in bAVM prevalence in male vs. female patients(Hofmeister et al. 2000; Lunsford et al. 2008; Brinjikji et al. 2017; Yang et al. 2019; Bameri et al. 2021). For murine experiments, postnatal animals that had died were excluded from vessel analysis (as they could not be reasonably perfused, for example), otherwise all animals were included in each mouse study.

#### Postnatal Induction of CreER activity

Tamoxifen (Sigma-Aldrich, cat#T5648) was equilibrated to room temperature, then reconstituted in warm (37°C) 100% ethanol at 100 mg/mL. Once in solution, this stock was then diluted 1:10 in warm sesame oil (Sigma) to a concentration of 10 mg/mL and single use, working, aliquots were frozen at -20°C and thawed at 37°C prior to injection. The 10 mg/mL stock solution was diluted 1:1 in sesame oil (for a final concentration of 5 mg/mL) and 40 µL of tamoxifen (thus, approximately 20 µg total per mouse) was administered by subcutaneous (s.q.) injection at postnatal day 1 using a 31G, 8mm insulin syringe (BD, cat#328438).

#### Postnatal CNS vascular-induction of KRAS activity

*Kras^G12D/WT^*; *R26^RFP/RFP^* or *Kras^G12C/WT^*; *R26^RFP/RFP^* females were crossed with males carrying a CNS-specific, endothelial-restricted CreER transgene, *Slco1c1(BAC)CreER* (*Slco1c1-CreER*; *Kras^WT/WT^*; *R26^WT/WT^*). All resulting offspring were given tamoxifen by s.q. injection at P1 (as outlined above).

#### In Vivo Inhibitor Treatment

Before initiation of the treatment, mice were randomized to receive Mirdametinib (Selleckchem, PD0325901) or vehicle. PD0325901 was dissolved in DMSO (50 mg/ml stock), and then diluted in water containing 0.05% hydroxypropyl-methycellulose and 0.02% Tween 80 with a final concentration of 2.5 mg/ml PD0325901. Aliquots were stored at -20C. Mirdametinib (PD0325901) or vehicle were given to mice daily by oral gavage (25 mg/kg) for 4 weeks (6 consecutive days/week).

#### Vascupaint and micro-CT imaging

Mice were anesthetized using isoflurane until they were unresponsive to noxious stimuli. Afterward, their chest was opened, rib cage reflected, and right atrium opened. Then, they were transcardially perfused through the left ventricle using a blunted 25-gauge syringe (BD PrecisionGlide, #305122) with 8 mL warm 1x PBS with heparin (Mckesson, #63739092025) (20 U/mL), followed by 8 mL 10% neutral buffered formalin (Leica, #3800598), and 1 mL Vascupaint^TM^ (MediLumine Inc, MDL-121). Vascupaint was prepared as follows: 1 mL silicone, 2 mL diluent, and 40 μL catalyst. After a successful perfusion, mice were left head down at 4°C. The following day, after the Vascupaint^TM^ solution had solidified, the brain was carefully resected from the skull, making efforts to prevent damage or loss of superficial vessels. Brains were then imaged using a Stemi stereomicroscope (Zeiss) and scored for hemorrhage as determined by evidence of blood pooling on the surface of the brain. After imaging, brains were embedded in 1% low melt agarose in ddH_2_O in a 5 mL plastic vial (Axygen Scientific, ST-5ML) to stop sample movement during imaging and to prevent shrinkage due to dehydration. Micro-CT images of the brains were obtained using a SkyScan 1272 (in the Optical and Intravital Microscopy core at BCM) or a SkyScan 1276 CMOS EDITION (in the Micro CT Imaging Facility at the McGovern Medical School at UTHealth). Images obtained on the SkyScan 1272 were collected using the following parameters: source voltage and current (60 kV and 166 μA), imaging voxel size was 10 µm with 0.25 µm AI filter, step projection set to 0.4 degree rotation, with a full 360 degree rotation (i.e. 1800 projection). The distance of X-ray to object was set to 174.60 mm. For imaging using the SkyScan 1276, scans were performed with the parameters: source voltage and current (60 kV and 166 μA), imaging voxel size was 10 µm with 0.25 µm AI filter. The distance of X-ray to object was set to 199.98 mm. Acquired images were then reconstructed using NRecon Reconstruction software (Bruker Micro CT, Kontich, Belgium) and then rendered into quantifiable 3D datasets in CT-Vox volume rendering software (Bruker Micro CT, Kontich, Belgium). The brains with an obvious venous or arterial dilation and shunt, visible in different locations of the brain after micro-CT imaging, were scored as positive for an AVM.

#### Visualization and Analysis of Brain Vasculature in CTAn

Image processing for quantitative analysis was performed using NRecon and CTAn software (Bruker). First, the images were reconstructed in NRecon (Bruker Micro CT, Kontich, Belgium) (output ranges from 0.0 to 0.1) and saved as BMP files. Second, region of interest [ROI] image series were loaded onto CTAn (Bruker Micro CT, Kontich, Belgium). Images were processed to create a stack of binary images and to eliminate background pixels (voxels in 3D) (Thresholding: mode, Global; lower grey threshold, 50; upper grey threshold, 255 / Despeckle: remove white speckles; 3D space; less than 100 voxels). Refined 2D stack image series were processed using CTAn 3D analysis to obtain vessel morphology parameters.

#### Visualization and Analysis of Brain Vasculature in VesselVio^TM^

VesselVio^TM^ software (Version 1.1.2, Bumgarner, 2022) was used to facilitate visualization and analysis of reconstructed brain images, yielding data on parameters such as vessel radius size, tortuosity, surface area, and vessel volume. CT image sequences were obtained using a Bruker 1272 or 1276 at 10 μm resolution and then the reconstructed image dataset was imported into the Fiji image analysis software(Schindelin et al. 2012). Subsequently, the image sequences underwent processing using the “median 3D filter” function. Next, image thresholding was performed using the Otsu Red setting. Particular attention was paid to adjusting the threshold to maintain a clear visualization of the vessels, using the threshold sliders. The processed image sequence was then saved using Fiji’s NIfTI (Neuroimaging Informatics Technology Initiative) plugin, resulting in a .nii file format file that is compatible with VesselVio^TM^. Subsequently, the .nii file was uploaded to VesselVio^TM^ using the “visualize” tab, and the resolution of the initial CT scan entered (which in our case was 10.0 μm). Then the rendering quality was set to “high” and the “scaled” option was selected for accurate visualization of the brain vasculature. For analysis, the “analyze” function was used by loading the .nii file of interest, yielding a folder with an excel spreadsheet of all the data extracted from the reconstructed brain. This data was then imported into PRISM software (Graphpad) for further analysis. Pairwise analyses for parametric tests were performed using Student’s t-test.

#### Visualization and Analysis of Brain Vasculature in Vesselucida360^TM^ and VesselucidaExplorer^TM^

Regions of interest [ROI] from the brain were visualized using Vesselucida360^TM^ (MBF Bioscience, Vermont, USA). Images were loaded in a TIFF format as a stack; with X, Y, and Z parameters set to 10.0 voxels. The sequence was then saved as an MBF JPEG file (.jpx). In the 3D environment, vessel tracing and 3D reconstruction was performed manually. The tracing option selected was voxel scooping, and appropriate seeds were selected to properly visualize the vasculature in detail. Seeds were refined and filtered and then saved in the 3D environment as a .xml file. Analysis of visualized tracing was completed in VesselucidaExplorer^TM^ (MBF Bioscience, Vermont, USA). The parameters quantified by VesselucidaExplorer^TM^ were vessel volume, surface area, total length, and the number of branching nodes. The extracted data were imported into PRISM software (Graphpad) for further analyses. Pairwise analyses for parametric tests were performed using Student’s t-test.

For comparing data between 2 month and 7 month old animals, each data point was calculated as follows: Percent change = ((Mutant ROI vessel length at 2 months – Average control ROI vessel length at 2 months)/Average control ROI vessel length at 2 months)*100)). The same formula was applied for 7-month-old samples.

#### Analysis of the Murine Retinal Vasculature

2-month-old adult mice, under anesthesia, were retro-orbitally injected using a 31-gauge insulin syringe (BD, 328438) with 50 µL of far red (649 nm) fluorescently conjugated lectin (*Lycopersicon esculentum*) (Vector Labs, DL-1178-1) to label the lumenal endothelium of patent vessels, and the dye was allowed to circulate for 5 minutes. Animals were then anesthetized using inhaled isoflurane until a surgical plane of anesthesia was reached. Next, their chest was opened, rib cage reflected, and they were transcardially perfused through the left ventricle using a 31-gauge insulin syringe with 100 μL of fluorescently labeled far red lectin (649). Then, the right atrium was opened, and the left ventricle perfused using a 27-gauge syringe with 10 mL of warm 1x PBS followed by 10 mL of 4% Paraformaldehyde (PFA) in 1x PBS. Eyes were then enucleated and placed in 4% PFA at 4°C with gentle agitation for 1 hour and then washed 3 times with 1x PBS. The retinas were then dissected out and once free were partially cut into 4 quadrants (i.e. leaflets) to allow for flat mounting. Isolated retinas were then blocked and permeabilized for 1 hour with gentle shaking at 4°C in retina blocking buffer (0.2% BSA / 0.3% Triton-X100 / 1x PBS). Retinas were then incubated overnight with gentle shaking at 4°C with an anti-actin, α-Smooth Muscle (αSMA) FITC-conjugated antibody (Sigma, F3777) [1:100] prepared in retina blocking buffer. The next morning, retinas were washed 4X, 10-minutes per wash, in PBS + 0.3% Triton (PBSTx) and then mounted on glass slides and coverslipped using Fluoromount-G mounting medium (SouthernBiotech, 0100-01).

To determine whether there were defects in arteriovenous patterning, wholemount images were obtained using a Zeiss Axio Zoom.V16 fluorescent stereoscope at a 110X magnification, with 9 tiled images collected (10% overlap) using the automated stage and the tiling function. Images were acquired in the green and far-red channel (2.3X objective, Plan Neofluar Z 2.3X, 500 ms exposure, total magnification of 110 and image size of 2752 x 2028 px). Absence or presence of AVMs were classified by visualization of direct connection between arteries (SMA) and veins (lectin).

To quantify vascular area, branching, and tortuosity, a single 20X image was taken of the central plexus for each quadrant of a retina (i.e. 4 images per retina) using a Zeiss LSM 780 confocal microscope with a 20X objective lens (1,024 x 1,024 pixels) at 15% laser power (561 nm). Effective NA of 0.8, pinhole 1.47 (au), line time of 30.00 μs, unidirectional scan, averaging of 1, -2° rotation, and 1.89 s scan time. Images were processed using Angiotool software (Zudaire et al. 2011) within each individual 20X image and averaged across the 4 quadrants for a single averaged measurement of vascular branching per biological sample. This was performed from 1 retina per animal for both control (n=8) and experimental (n=8) groups. Data was graphed using GraphPad Prism software and shown as an average ± SEM. Pairwise analyses for parametric tests were performed using Student’s t-test.

#### Representative Images

For all images included in the manuscript, representative examples that reflect the typical phenotype were chosen. Quantification of phenotypes was performed where possible, and the chosen images reflect the median of the quantification. In some cases, several images were included when there was a range of possible phenotypes.

#### Neurobehavioral Evaluation

All animal studies conducted were approved by the Institutional Animal Care and Use Committee at Baylor College of Medicine, the University of Virginia School of Medicine, and UT Health Sciences Center at Houston. Mice were maintained on a 12:12 hour light cycle with standard mouse feed (Pico Lab Rodent Diet, #5053, Lab Diet, St. Louis, MO, USA) and water *ad libitum*. All tests were conducted between 9:00 AM and 6:00 PM. The behavior battery comprises the following assays: Open field, parallel rod footslip, catwalk, novel object recognition and conditioned fear. For each assay, mice were habituated to the test room for at least 30 min prior to testing. All behavior equipment used in our study were cleaned with 35% isopropanol between tests. Experimenters were blinded to the genotypes during testing and data scoring.

#### Locomotor Activity

Locomotor activity was assessed using the VersaMax Animal Activity Monitoring System (Omnitech Electronics, Columbus, OH). The test arena comprised of a clear acrylic (length: 40 cm; width: 40 cm; height: 30 cm) chamber placed in an enclosure containing panels of photo beams. The test animals were placed in the center of the arena under dim lighting conditions of 150-200 lux and allowed to explore the chamber for 30 min. The animal’s movement was assessed using photo beam breaks and total distance travelled and vertical activity were recorded using the VersaMax software version 4.2.

#### Parallel Rod Footslip

Fine motor coordination was assessed using the parallel rod floor test apparatus (Stoelting, Wood Dale, IL, USA). The animals were placed in a plexiglass box with a floor comprising of metal rods raised about 1cm above a steel base plate. The subject was placed in the test chamber and allowed to freely move for a period of 10 minutes. Every time the subject slipped, and the animal’s paw fell through the parallel rods and touched the base plate, a foot fall was recorded (ANY-maze software, Stoelting, Wood Dale, IL, USA). Locomotor activity was recorded using the overhead video camera and quantified using the ANY-maze software. Foot slips were normalized to total distance traveled by the subjects during the test period.

#### Spontaneous Gait Analysis (Catwalk XT)

Mice were allowed to freely walk on a transparent surface that was covered, creating a long dimly lit corridor with a high-speed digital video camera mounted below the surface. Each subject was allowed to explore the compartment until 3 compliant crossings across the view of the camera were achieved. Using the high-speed video camera, illuminated paw prints were captured and gait parameters were generated using Noldus CatWalk XT software. If an animal chose not to participate in the test they were removed from the apparatus and retested later.

#### Novel Object Recognition

Our apparatus comprised of a rectangular plastic arena (45 cm long x 24 cm wide x 20 cm high) surrounded on three sides by a white opaque screen. Three angled mirrors were placed behind the cage to enable the observer to see the back side of the arena. On day 1, animals were allowed to freely explore the test arena without the objects (habituation phase) for 5 minutes. Immediately following the habituation phase the animals were placed in the test arena containing 2 identical LEGO objects and allowed to explore for 5 minutes. On day 2, following the habituation phase similar to day 1, animals were placed in the test arena with one familiar object from the previous day and one novel object and allowed to explore for 5 minutes. The time spent at the objects was scored in real time by observers blinded to genotypes. An overhead camera set up was used to record the entire experiment and was connected to a video-assisted tracking software package (ANY-maze Behavioral tracking software, Stoelting Co). The novel object recognition index was calculated as follows: [time spent at the novel object/total time spent at both object] x 100. This assay was performed under dim lighting condition of 150 lux with background white noise 60 dB.

#### In Situ Hybridization (smFISH) and Confocal Imaging

Adult brains from male C57BL/6 mice were collected and immersion fixed in freshly made 4% PFA/1x PBS/0.1% Tween-20 overnight at 4°C with gentle agitation. The following day, tissues were extensively washed then mounted in 1% agarose and sectioned at 35 μm thickness using a compresstome (World Precisionary Instruments, VF-300-Z). Sections were then mounted on Superfrost Plus Microscope Slides (Fisher Scientific, #22-037-246) and dried at 50°C for 2 hours. Slides were then washed with 1X PBS for 5 minutes before being dried for an hour at 60°C. An ImmEdge Hydrophobic Barrier Pen (Advanced Cell Diagnostics, Cat No #310018) was used to establish a hydrophobic barrier around each of the samples prior to fixing the samples in 10% formalin for 15 minutes at 4°C.The samples were then serially dehydrated through a gradient of 50%, 70%, and 100% ethanol at room temperature. Antigen retrieval was then performed using RNA Target Retrieval Reagent (Advanced Cell Diagnostics, Cat No. PN 322000) followed by a secondary dehydration step consisting of a 5 minute incubation at room temperature in 100% ethanol. After complete dehydration, the samples were then incubated with Protease III for 30 min at 40°C using the HybEZ oven (Advanced Cell Diagnostics, Cat No. 321720) before washing 5 times in ddH_2_O, 1 minute per wash. Sections were then incubated in the *Kras* RNA Scope probe (Advanced Cell Diagnostics, Cat No #412491, ∼5-7 drops per section) for 2 hours at 40°C. The slides were then washed twice in 1x wash buffer (Advanced Cell Diagnostics, Cat No. 310091) for 5 min per wash. Amplification and detection steps were performed using the RNAScope Multiplex Fluorescent Reagent kit v2 (Advanced Cell Diagnostics, Cat. No. 323100). Sections were incubated with Amp1 for 30 min at 40°C and then washed 2 times in 1x wash buffer (Advanced Cell Diagnostics) for 5 min. This process was then repeated for the Amp2 and Amp3 steps. Samples were then incubated in Horseradish Peroxidase for Channel 1 (HRPC1, Advanced Cell Diagnostic) for 15 min at 40°C, before washing two times in 1x wash buffer for 10 minutes at room temperature. The samples were then incubated in HRP blocking buffer for 30 min at 40°C, followed by two washes in 1x wash buffer for 10 minutes per wash. Samples were then incubated in freshly prepared blocking buffer (10% goat serum/1X PBS/0.2% TritonX-100) for an hour at room temperature before incubating in Alexa Fluor 488 conjugated anti-Claudin-5 (ThermoFisher #4C3C2, #352588), diluted in blocking reagent (1:100 dilution) overnight at 4°C.

The following day, samples were washed in 1x wash buffer (Advanced Cell Diagnostics) prior to stanining with DAPI (ACD Biotechne, Cat. No. 320858) to label all nuclei. Samples were then mounted with Prolong Gold Antifade Mounting Medium (ThermoFisher, Cat. No. P36930). Each assay was conducted alongside a positive control treated with PPIB (ACD Biotechne, Cat. No. 540651) and a negative control slide treated with DapB (ACD Biotechne, Cat. No. 707351).

Entire midsagittal sections of the adult mouse brain were imaged by confocal microscopy with a step size of 2.5 μm, at a depth of 10 slices in the z-plane, using an LSM780 confocal laser scanning microscope (Zeiss) and images were collected with 20% overlap to enable tiled reconstruction of the entire brain using a 10x objective (NA=0.45), with laser power of 10.0% (405 nm), 30.0% (488 nm), 8.0% (560 nm), and a pinhole of 24.6 μm. Confocal stacks were compressed to a single Maximum Intensity Projection (MIP) and the tiled MIP images were stitched together in Zen (black edition) software (Zeiss).

For quantification of *Kras* transcripts in each region of the brain, confocal images were collected at a step size of 1 μm, across a depth of 22 slices, using an LSM780 confocal laser scanning microscope (Zeiss) with a 40x (NA=1.4) oil objective, and laser power of 2.0% (405 nm), 22.0% (488 nm), 2.0% (560 nm), and a pinhole of 51 μm. A total of three confocal stacks were taken from each region per brain. The confocal stacks were then compressed to a Maximum Intensity Projection (MIP) using the ZenBlack imaging software (Zeiss) before being exported for quantitative analysis (as described below).

#### Quantitative Analysis of smFISH

To quantify *Kras* transcript levels within the cerebrovasculature in various regions of the adult mouse brain, quantitative analysis of smFISH signal was conducted as described(Secci et al. 2023). Briefly, three random maximum intensity projection (MIP) images (each measuring 1024 by 1024 pixels) of sections stained with CLDN5 (to label all endothelial cells), DAPI (to label nuclei), and *Kras* smFISH (to detect *Kras* transcripts) were collected from six regions of the adult murine brain (olfactory bulb, cortex, cerebellum, hippocampus, thalamus, and hypothalamus). Images were processed using QuPath(v.0.5.1-x64), an open source morphometric analysis package(Bankhead et al. 2017). CLDN5 immunoreactivity was used to identify endothelial cells and to facilitate manual tracing of the region of interest [ROI]. Endothelial cells residing within the perimeter of vessels greater than 10 μm in diameter were used for quantification analysis of smFISH. Using this criteria, *Kras*-positive mRNA puncta in CLDN5^+^-endothelial cells were quantified using QuPath’s cellular and subcellular detection algorithm. QuPath cell detection was set to identify cells ranging between 10-200 μm^2^, with a surrounding expansion of 3 μm per cell. mRNA puncta were scored as positive using a subcellular detection threshold of 500 and by areas ≥0.5 and ≤19 μm^2^. The resulting data were exported to Excel, and the average intensity and the average number of puncta per CLDN5+ cell was calculated. Data were then graphed using PRISM software (Graphpad) and are shown as the mean +/- s.e.m. Statistical significance was determined through a multi-way ANOVA and Tukey’s post-hoc analysis, with p<0.05 considered statistically significant.

#### Statistical Analyses

Unless otherwise indicated, experiments were performed a minimum of 3 independent times. The number of replicates is included in figures or figure legends and exact p-values are indicated in the figures. Normal distribution of data was assessed visually by QQ plot and statistically by the Shapiro-Wilk test (p>0.05). Parametric tests were utilized for data that featured a normal distribution. Pairwise analyses for parametric tests were performed using Student’s t-test, while comparisons of 3 or more groups were conducted using a one-way ANOVA and Tukey’s post-hoc analysis. All statistical data was processed, and graphs were generated, using Prism 9 (Graphpad).

## ACKNOWLEDGEMENTS

The authors thank Jorge Castro and Maci Heal at MBF Bioscience for guidance with Vesselucida software and Ali Bahadur at Bruker for access to, and assistance with, NRECON, CTAN and CT-VOX software. Research reported in this publication was supported by the Eunice Kennedy Shriver National Institute of Child Health & Human Development of the National Institutes of Health under Award Number P50HD103555 for use of the Preclinical Outcomes Measure Core facilities. Micro-CT imaging was supported by NIH S10OD030336 and performed through the Micro-CT Imaging Facility at the McGovern Medical School at UTHealth and in the Optical and Vital Microscopy Core facility at Baylor College of Medicine. This work was supported by grants from the Canadian Institutes of Health Research (CIHR PJT155922), the U.S. Department of Defense (W81XWH-18-1-0351) and the National Institutes of Health (1R01HL159159-01A1) to J.D.W. and J.E.F; from the American Heart Association to CFS (AHA award - 916015); and by generous seed funding from the Cardiovascular Research Institute at Baylor College of Medicine and the Joe Niekro Foundation to J.D.W., S.G.M. III, and I.I and from the University of Virginia School of Medicine to J.D.W.

## AUTHOR CONTRIBUTIONS

Author Contributions: JDW conceptualized the study; JDW and CFS designed experiments; CFS, OAH, JCIII and EKA conducted the majority of experiments with help from GL and SV; AR and PK performed the retrospective case study; S.M. and I.I. provided deidentified patient data; CFS, OAH, SM, SV, JEF and JDW analyzed the data; CFS and JDW wrote the original draft of the manuscript; JDW, JEF, SM, and CFS secured funding; all authors edited and approved the manuscript.

## SUPPLEMENTAL FIGURE LEGENDS

**Supplemental Figure 1.**
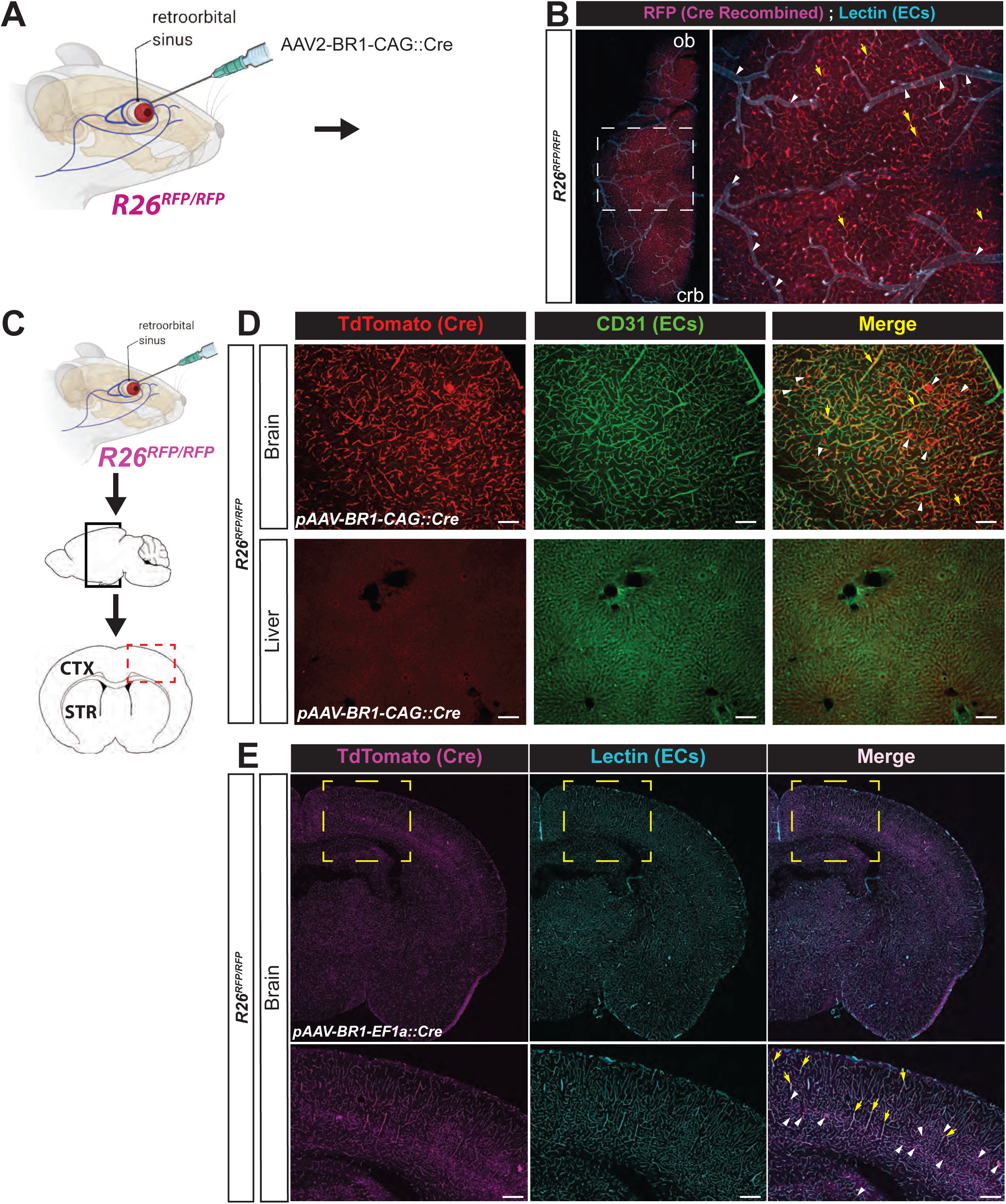
AAV-BR1 has broad tropism within the central nervous system. (**A**) Experimental overview of the retroorbital delivery of AAV-BR1-CAG::Cre in mice. (**B**) Representative image of an the brain from an adult *Rosa26^lsl-RFP^* (*R26^RFP^*) mouse transduced with AAV-BR1-CAG::Cre following transcardiac perfusion with far red fluorescent lectin (cyan) to label all vessels. Ob=olfactory bulb, crb=cerebellum. The white, boxed area in the left panel is magnified on the right. Yellow arrows denote RFP labeling of small capillary vessels, while white carets denote the absence of RFP signal in large diameter vessels. (**C**) Experimental overview for delivery of AAV-BR1-CAG::Cre and isolation of coronal sections from the adult mouse brain. (**D**) Coronal sections from R26^RFP^ mice transduced with AAV-BR1-CAG::Cre. Immunostaining for endothelial cells (ECs) using antibodies against CD31, and visualization of Cre activity via the TdTomato Cre reporter, confirm the brain specific tropism of the virus (note the absence of RFP signal in the liver). Of note, the top row, far right column shows extensive recombination of the RFP reporter in non-endothelial cells (CD31^-^), highlighted by white carets, as cells with a neuronal-like morphology are labelled with TdTomato. Yellow arrows denote RFP-positive endothelial cells (RFP^+^, CD31^+^). Scale bar = 100 μm. (**E**) Substituting the synthetic CAG promoter for the less robust *EF1α* promoter somewhat mitigated transduction of non-endothelial cells. Confocal micrographs from sections of the adult murine brain from *R26^RFP^* mice transduced with AAV-BR1-EF1a::Cre, then later perfused with fluorescent lectin to label the endothelium, are shown. The boxed area within the cortex in the top row are magnified and shown in the bottom row. Yellow arrows denote vessels labelled by RFP, while white arrows denote non-endothelial cells transduced by AAV-BR1-EF1a-Cre.

**Supplemental Figure 2.**
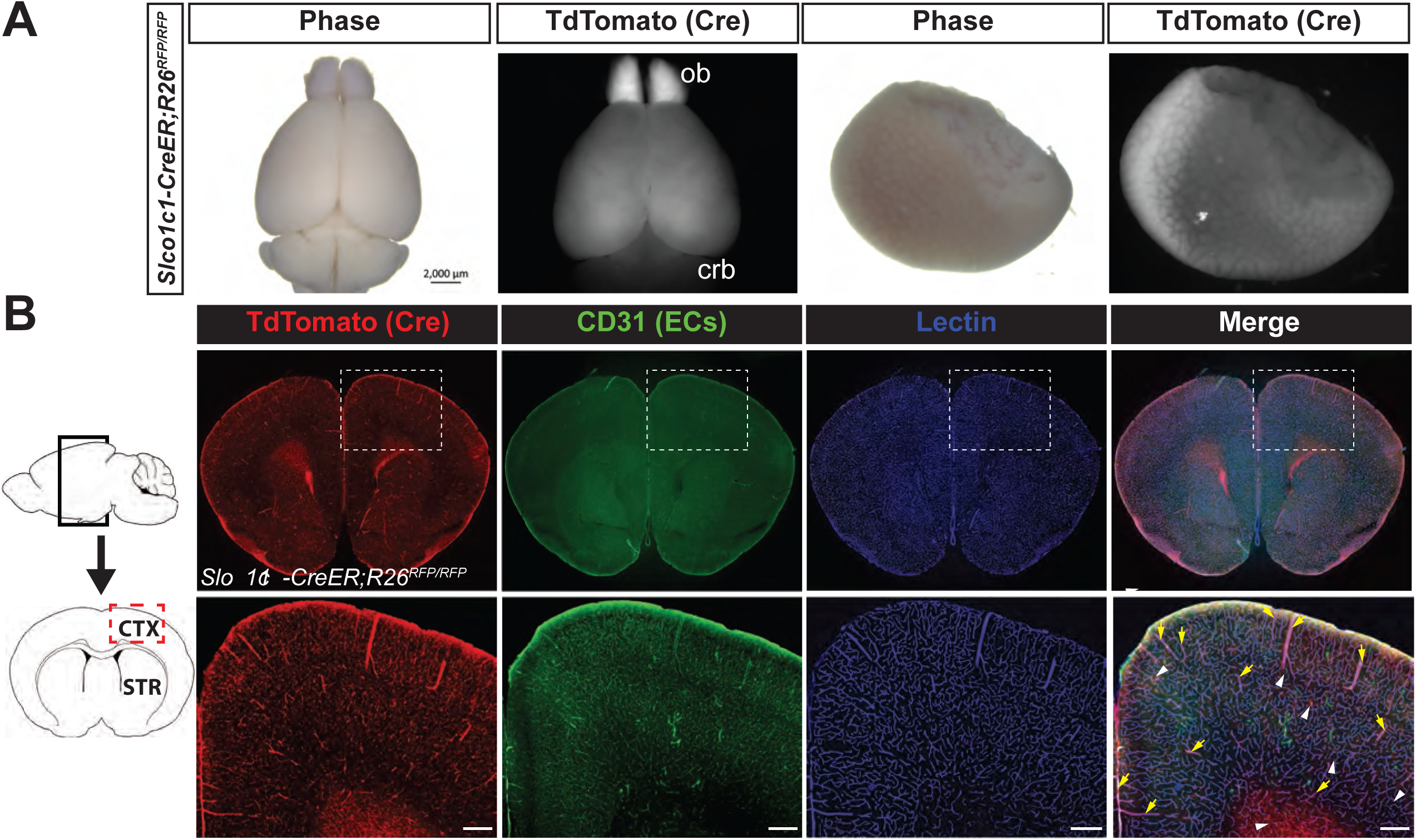
S*l*co1c1*-BAC-CreER* drives recombination through the postnatal murine CNS cerebrovascular endothelium. **(A)** Representative phase and epifluoresensce images of an adult murine brain [left] and liver [right], respectively, from a *Slco1c1-BAC-CreER*;*Rosa26^lsl-RFP^* (*Ai14*) mouse following administration of tamoxifen at P1. Ob=olfactory bulb, crb=cerebellum. Note the absence of RFP signal in the liver. (**B**) On the far left, a schematic shows the region where coronal sections were collected from for examining *Slco1c1-BAC-CreER* recombination in the adult murine brain. Cre activity is visualized by the TdTomato reporter [red], while the endothelium is visualized by staining for CD31 [green] and perfused vessels are labeled by lectin [blue]. The far right panel shows the merged image. The dashed, boxed regions in each panel of the top row are magnified and shown below. Note the extensive recombination in both large and small diameter vessels (yellow arrows), as well as sparse labeling of non-endothelial cells (white carets). Scale bar = 100 μm.

**Supplemental Figure 3.**
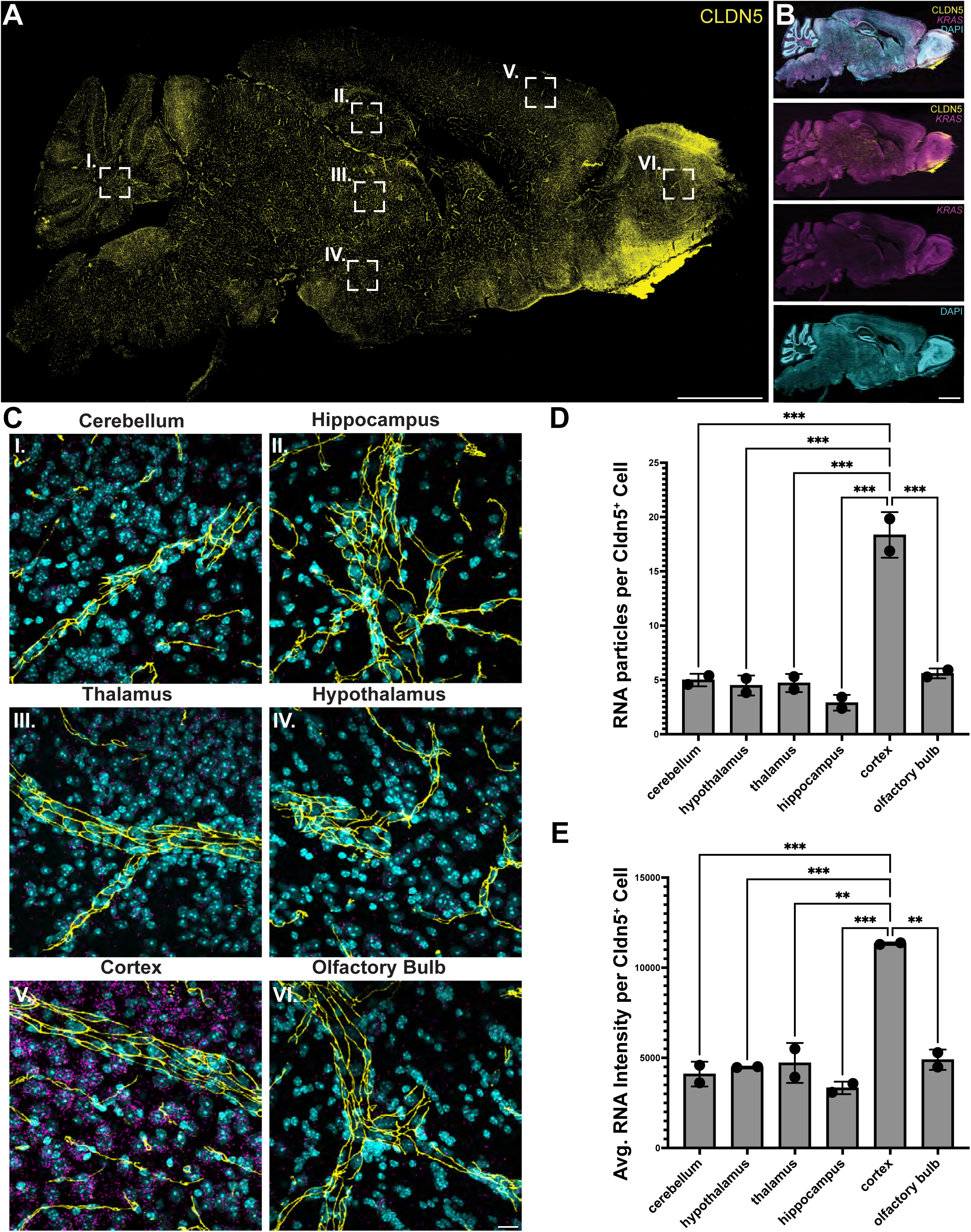
K*r*as transcripts are enriched in the adult murine cortex. **(A)** Representative image (sagittal view; cerebellum to the left, olfactory bulb to the right), of a section from an adult murine brain following immunostaining for the endothelial cell marker CLDN5 (yellow). Scale bar = 3 mm. (**B**) The far-right column shows images from the same processed for smFISH to detect *Kras* transcripts (magenta) and counter-stained with DAPI (cyan) to label all nuclei. Scale bar = 20 μm. CLDN5-positive endothelial cells in each region of interest (i through vii) were scored for the number, and intensity, of *Kras*-positive puncta. (**C**) Quantification of the number of *Kras* particles per CLDN5^+^ cell in each region. (**D**) Quantification of the intensity of *Kras* transcripts per CLDN5^+^ cell in each region. Data are presented as the mean +/- s.e.m. and are pooled from a total of 750 CLDN5^+^ cells, per region, per mouse (n=2 mice). * p≤0.05; ** p≤0.01; *** p≤0.001.

**Supplemental Figure 4.**
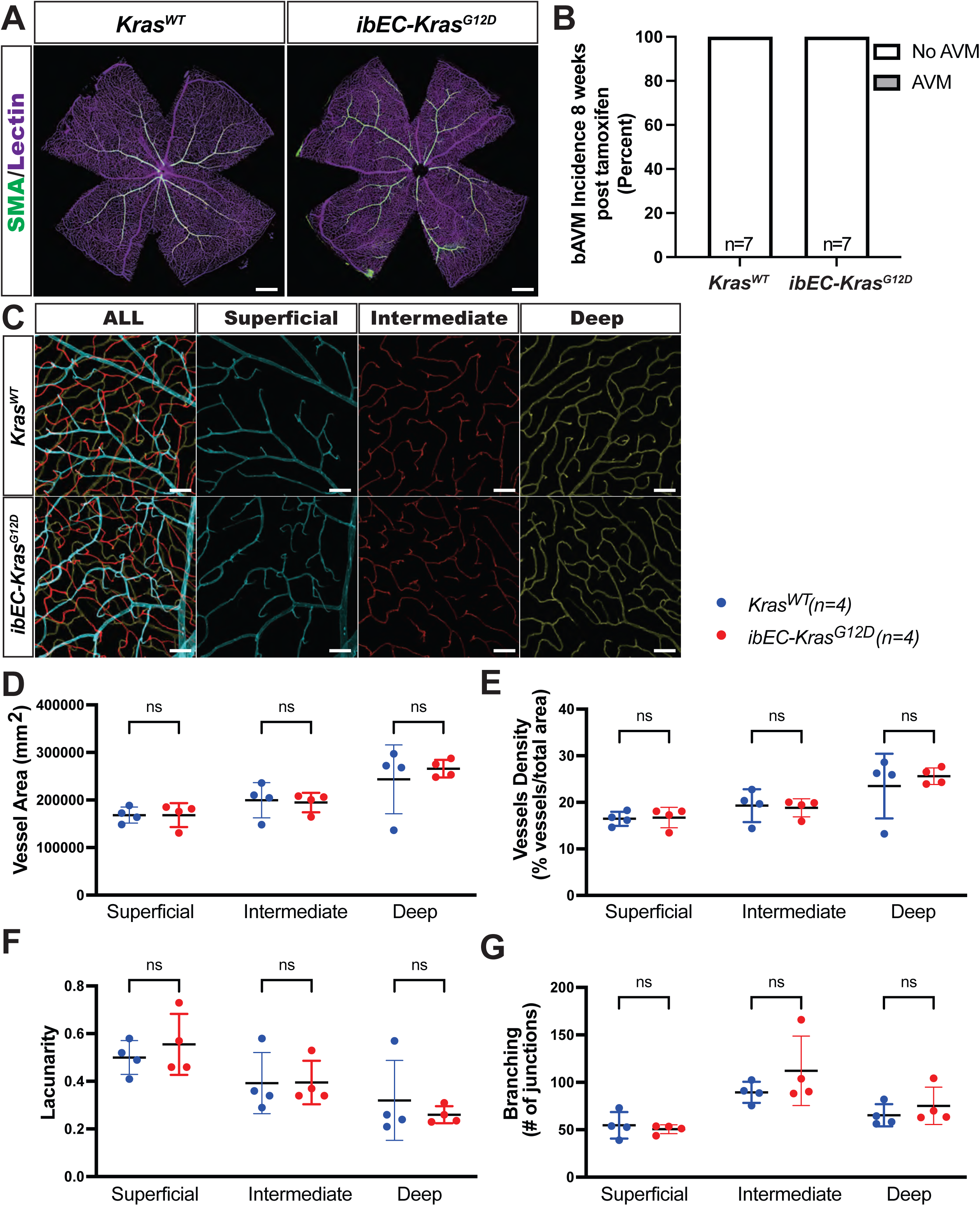
Endothelial-specific expression of mutant Kras does not lead to AVMs in the adult murine retina. (**A**) Whole-mount fluorescence microscopy images of retinal-flat mounts following perfusion with far red fluorescent red lectin to label vessels and immunostaining for smooth muscle actin (SMA) to label arteries fails to reveal arteriovenous malformations, or shunts, in 2-months-old *ibEC-Kras^G12D^* and *Kras^WT^* mice (n=7, each group). Scale bar = 500 μm. (**B**) Quantification of observed AVMs within the murine retinal vasculature at 8 weeks of age following tamoxifen injection at P1. Representative images of superficial, intermediate, and deep layers of the retinal vasculature in 2-months-old *ibEC-Kras^G12D^* and *Kras^WT^* mice. A color-coded depth projection (far left column) shows each vascular layer merged in a single maximal projection, while single layers show the superficial (cyan), intermediate [blue], and deep (yellow) layers imaged by confocal microscopy. Scale bar = 100 μm. (**C**) No significant differences in vessel area were detected between *Kras^WT/+^* (n=4) and *ibEC-Kras^G12D^* (n=4) mice in the superficial, intermediate or deep layer. Student’s T-test was conducted for each layer comparison. Superficial, p=0.9995; intermediate, p=8271; deep, p=0.5700. (**D**) Vessel density in the superficial, intermediate, and deep layer vasculature is indistinguishable between *Kras^WT^* and *ibEC-Kras^G12D^* mice. A Student’s T-test was conducted for each comparison. Superficial, p=0.8474; intermediate, p=0.9249; deep: p=0.5787. (**E**) Branching (cell junction number) is similar in each layer of the retinal vasculature between *Kras^WT^* and *ibEC-Kras^G12D^* mice. A Student’s T-test was conducted for each layer comparison. Superficial, p=0.5975; intermediate, p=0.2815; deep, p=0.4180. (**F**) Lacunarity is not significantly different in any of the three layers of the retinal vasculature between *Kras^WT^* and *ibEC-Kras^G12D^* mice. Student’s T-test was conducted for each layer comparison. Superficial, p=0.4795; intermediate, p=0.9756; deep, p=0.5106. (**G**) Branching (cell junction number) is similar in each layer of the retinal vasculature between *Kras^WT^* and *ibEC-Kras^G12D^* mice. Student’s T-test was conducted for each layer comparison. Superficial, p=0.5975; intermediate, p=0.2815; deep, p=0.4180.

**Supplemental Figure 5.**
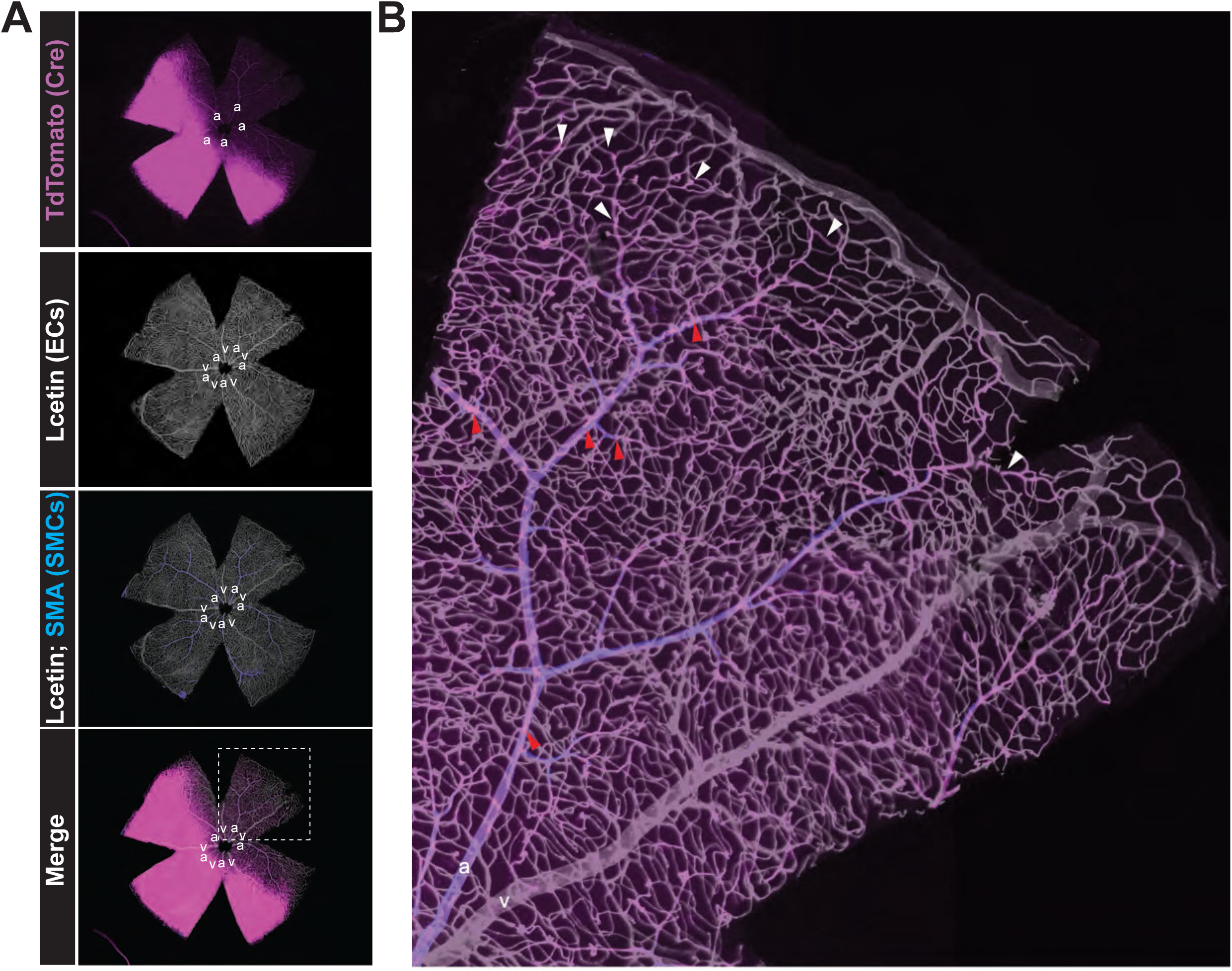
Slco1c1-CreER Drives Recombination in the Arterial and Capillary Endothelium of the Murine Retina. (**A**) Representative images of retinas from adult *Slco1c1-BAC-CreER*; *Ai14* mice following tamoxifen administration at P1 reveals extensive recombination in the arterial and capillary endothelium. Mice were perfused with fluorescent lectin shortly before euthanasia, and then retinas were processed for immunostaining using antibodies against the smooth muscle marker Smooth Muscle Actin (SMA). a=artery, v=vein. (B) The magnified image of the leaflet shows extensive Cre reporter signal (pseudocolored in magenta) in the arterial (blue) (denoted by red carets) and capillary endothelium (denoted by white carets), but not in the adjacent large diameter veins of the murine retina.

**Supplemental Figure 6.**
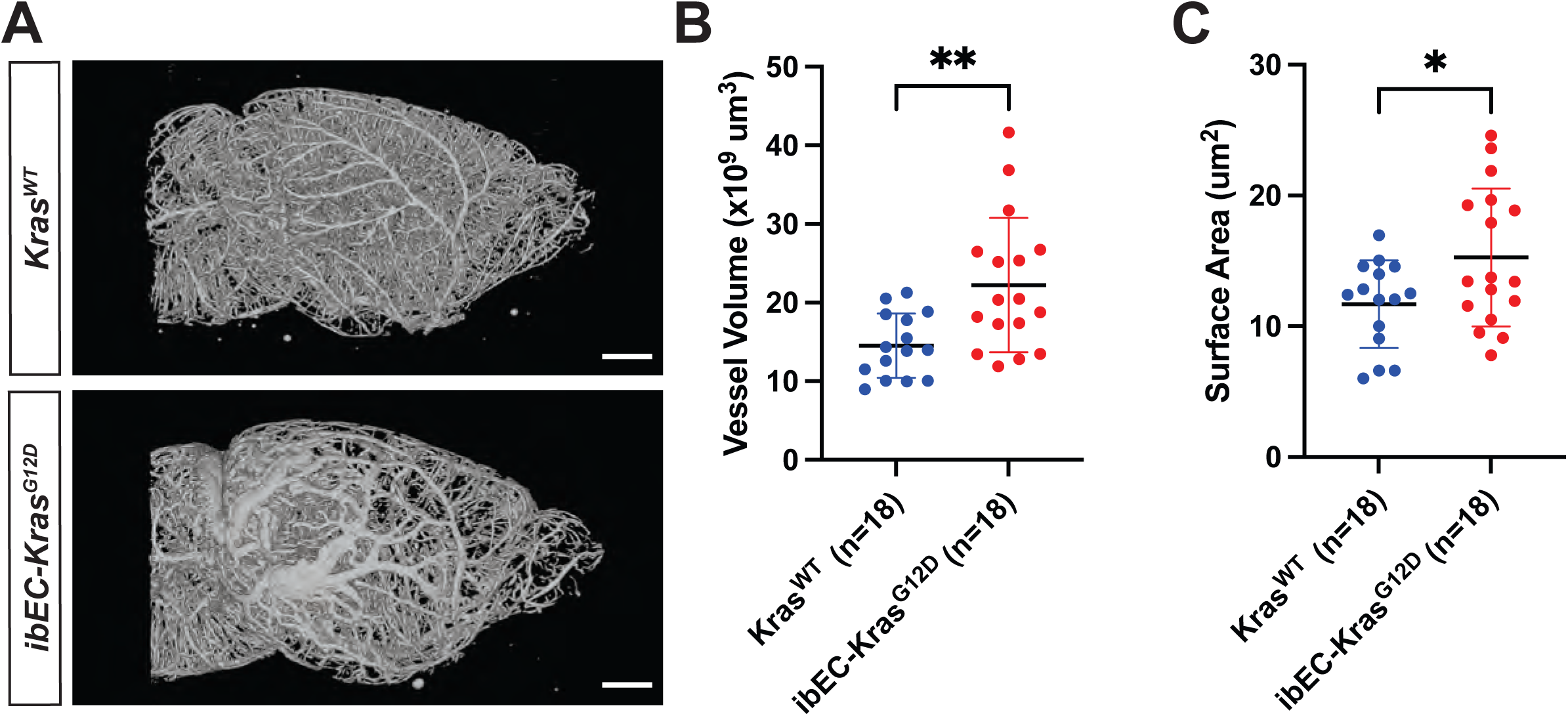
CT-AN reveals increased vessel volume and surface area throughout the cerebrovasculature in *ibEC-Kras^G12D^* mutants. (**A**) Representative sagittal views of the 2 month old murine cerebrovasculature following perfusion and micro CT image and vessel tracing and reconstruction using CT-AN software. Scale bar = 2000 μm. (**B**) Quantification reveals significantly increased vessel volume in *ibEC-Kras^G12D^* mice compared to *Kras^WT^* controls at 2 months of age (each dot represents one brain from a unique mouse, **p= 0.0034, student’s T-test). (**C**) Quantification of total surface area in control *Kras^WT^* and *ibEC-Kras^G12D^* mice (each dot represents one brain from a unique mouse, *p=0.0314, student’s T-test).

**Supplemental Figure 7.**
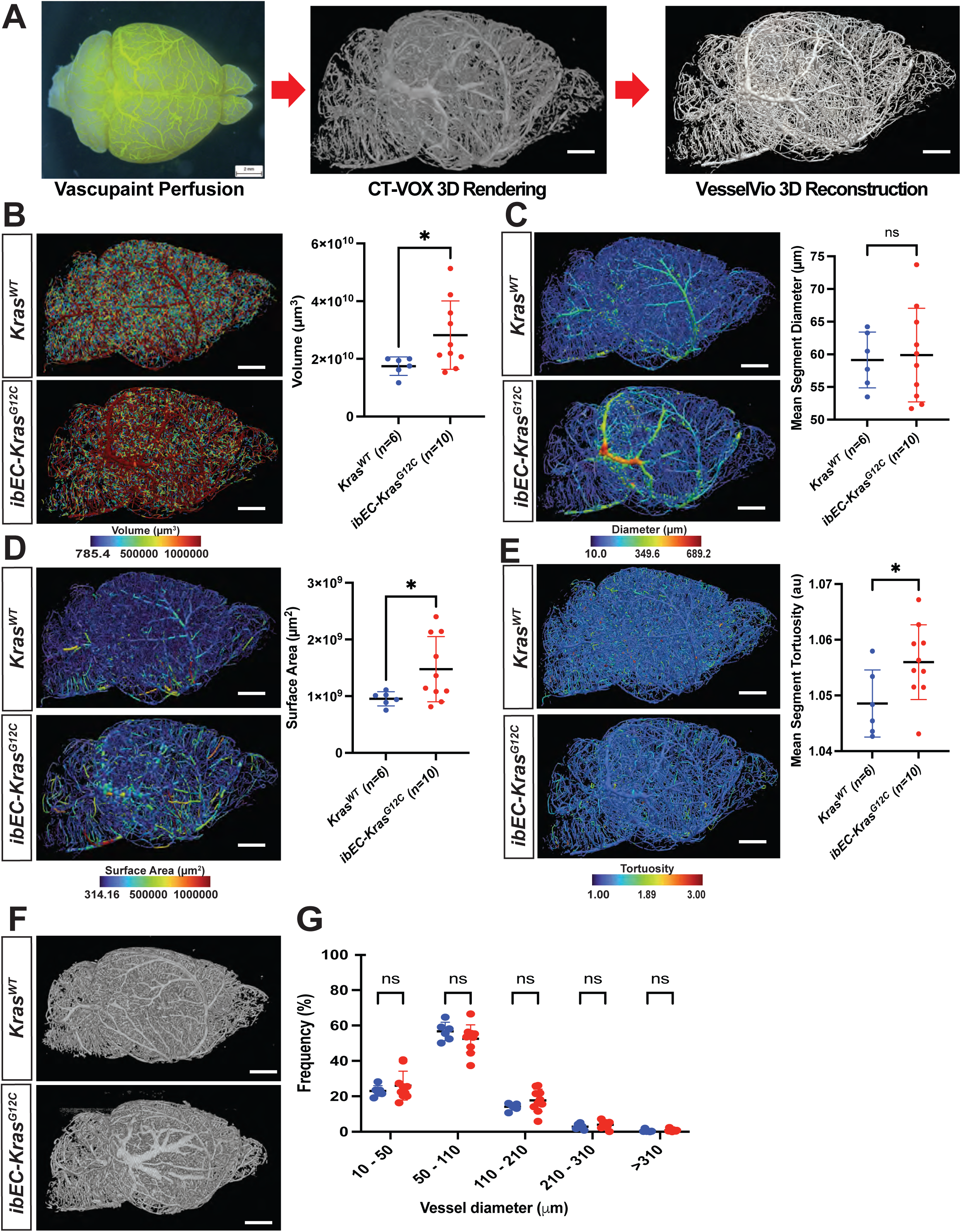
i*b*EC*-Kras^G12C^* mice feature increased vessel diameter, volume, and tortuosity across the entire cerebrovasculature. (**A**) Vessel reconstruction pipeline. The arterial circulation is labelled using Vascupaint contrast agent, followed by 3D micro CT imaging and CT-VOX 3D rendering. Vessel tracing and morphometric analysis are then performed using VesselVio software. (**B**) Sagittal view of representative heat maps from 2 month old adult mouse brains color-coded by vessel volume in (top left) control *Kras^WT^* (n=6) and (bottom left) *ibEC-Kras^G12C^* (n=10) mice. Right side, quantification of total vessel volume for each genotype (each dot represents one unique brain). Scale bar = 2000 μm. *p=0.0492 (Student’s T-test) (**C**) Sagittal view of representative heat maps from 2 month old adult mouse brains color coded by vessel diameter in (top left) control *Kras^WT^* (n=6) and (bottom left) *ibEC-Kras^G12C^* (n=10) mice. On the right side, quantification of mean segment diameter for each genotype (each dot represents one unique brain). Scale bar = 2000 μm. p=0.8191 (Student’s T-test). (**D**) Sagittal view of representative heat maps from 2 month old adult mouse brains color coded by vessel surface area in (top left) control *Kras^WT^* (n=6) and (bottom left) *ibEC-Kras^G12C^* (n=10) mice. On the right side, quantification of total vessel surface area for each genotype (each dot represents one unique brain). Scale bar = 2000 μm. *p=0.0482 (Student’s T-test). (**E**) Sagittal view of representative heat maps from 2 month old adult mouse brains color coded by mean segment tortuosity in (top left) control *Kras^WT^* (n=6) and (bottom left) *ibEC-Kras^G12C^* (n=10) mice. Right side, quantification of mean segment tortuosity for each genotype (each dot represents one unique brain). Scale bar = 2000 μm. *p=0.0437 (Student’s T-test). (**F**) Sagittal view of a vessel tracing using CT-AN for 3D reconstruction following micro CT imaging. Scale bar = 2000 μm. (**G**) Quantification of the frequency of various diameter vessels in control *Kras^WT^* and *ibEC-Kras^G12C^* mice (each dot indicates one brain from a unique 2-month-old mouse). Student’s T-test were conducted for each comparison. For comparison at 10 – 50 μm, p=0.442; 50 – 110 μm, p=0.0984; 110 – 210 μm, p=0.393; 210 – 210 μm, p=0.368; >310 μm, p=0.860.

**Supplemental Figure 8.**
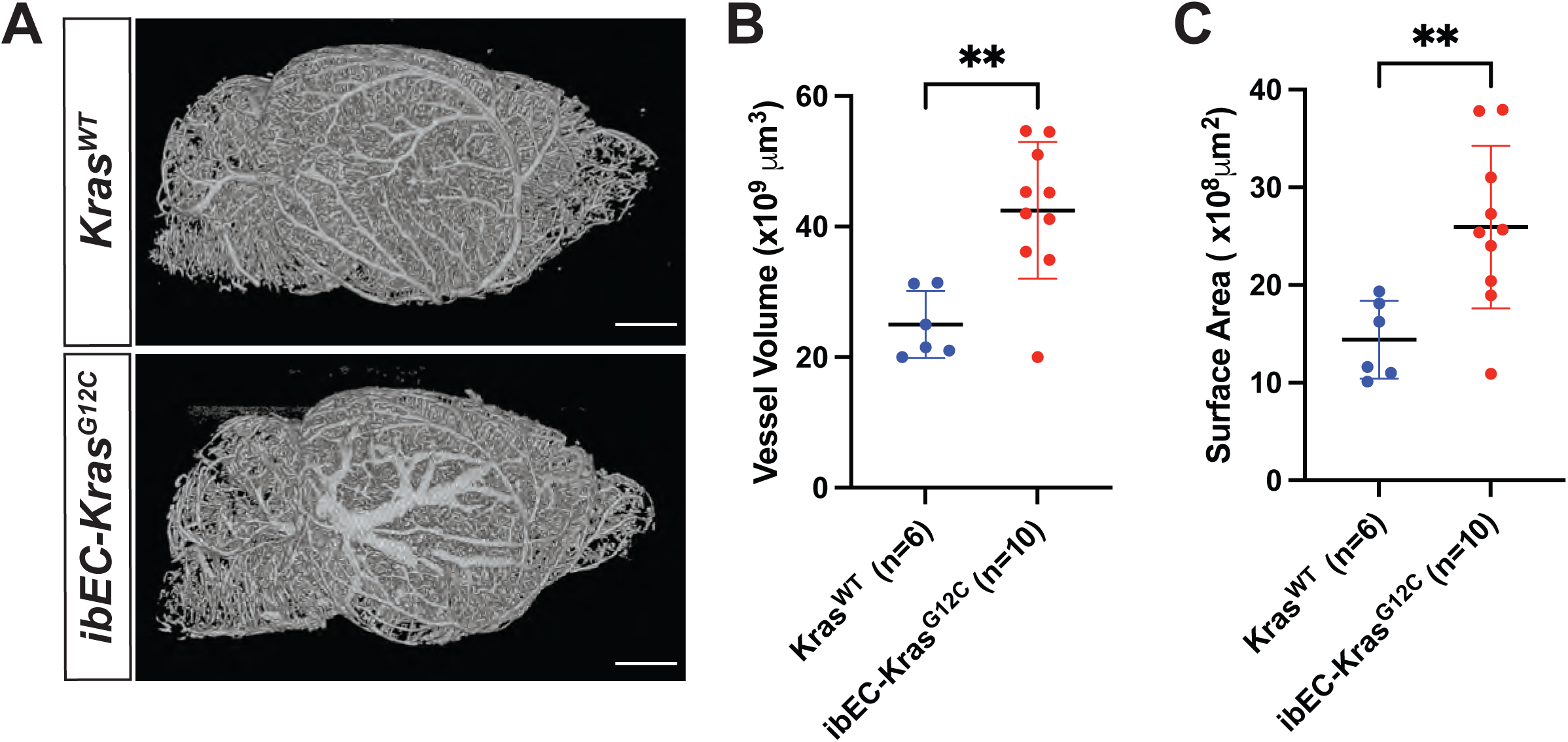
CT-AN reveals increased vessel volume and surface area throughout the cerebrovasculature in *ibEC-Kras^G12C^* mutants. (**A**) Representative sagittal views of the 2 month old murine cerebrovasculature following perfusion and micro-CT image and vessel tracing and reconstruction using CT-AN software. Scale bar = 2000 μm. (**B**) Quantification reveals significantly increased vessel volume in *ibEC-Kras^G12C^* mice compared to *Kras^WT^* controls at 2 months of age (each dot represents one brain from a unique mouse, **p= 0.0020 student’s T-test). (**C**) Quantification of total surface area in control *Kras^WT^* and *ibEC-Kras^G12C^* mice (each dot represents one brain from a unique mouse, **p=0.0070 student’s T-test).

**Supplemental Figure 9.**
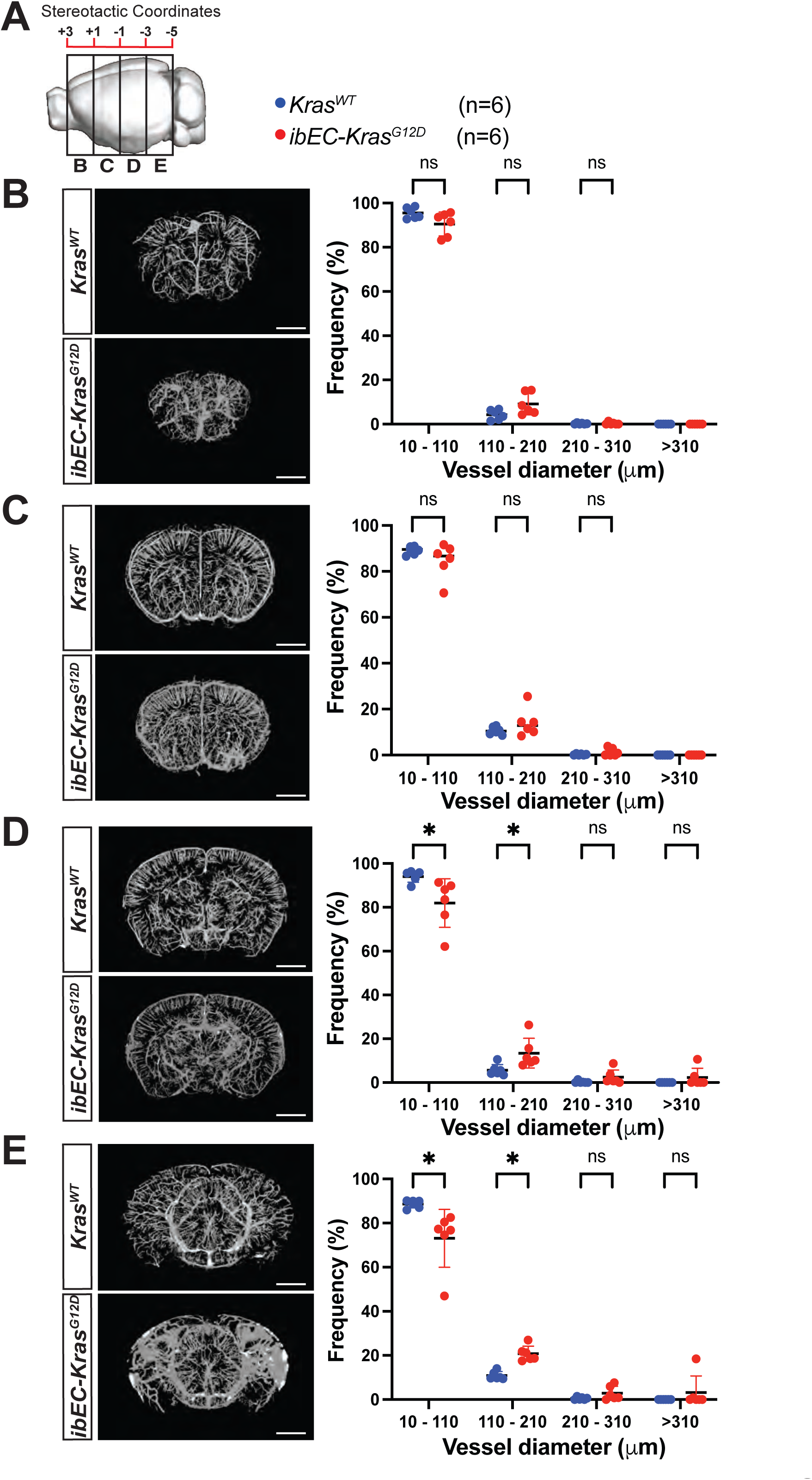
K*r*asG12D expression induces regional-specific changes in the cerebral vasculature. **(A)** A model illustrating the various 2,150 micron thick virtual coronal slab sections from micro CT used for the quantitative analysis with CT-AN shown in panels (B) through (E). (**B**) [ROI] A representative image of a 2,150 micron thick virtual coronal plane section located in the +3 - +1 region. (Right) No significant differences in vessel frequency were observed between *Kras^WT^* and *ibEC-Kras^G12D^* mice. (each dot represents a unique brain). (**C**) [ROI] A representative image of a 2,150 micron thick virtual coronal plane section located in the +1 - -1 region. (Right) No significant differences in vessel frequency were observed between *Kras^WT^* and *ibEC-Kras^G12D^* mice. (**D**) (Ivanova et al.)A representative image of a 2,150 micron thick virtual coronal plane section located in the -1 - -3 region. (Right) *ibEC-Kras^G12D^* mice featured a significantly decreased frequency of small vessels (10-110 μm) than *Kras^WT^* littermates. Student’s T-test was conducted for each layer comparison. For 10-110 μm: *p=0.0258, for 110-210 μm: *p=0.0263. (**E**) [ROI] A representative image of a 2150 micron thick virtual coronal plane section located in the -3 - -5 region. (Right) *ibEC-Kras^G12D^* mice showed significant differences in the frequency of small (10-110 μm)(*p=0.0165, Student’s T-test) and medium (110-210 μm)(*p=0.0000960), Student’s T-test) diameter vessels compared to *Kras^WT^* littermates.

**Supplemental Figure 10.**
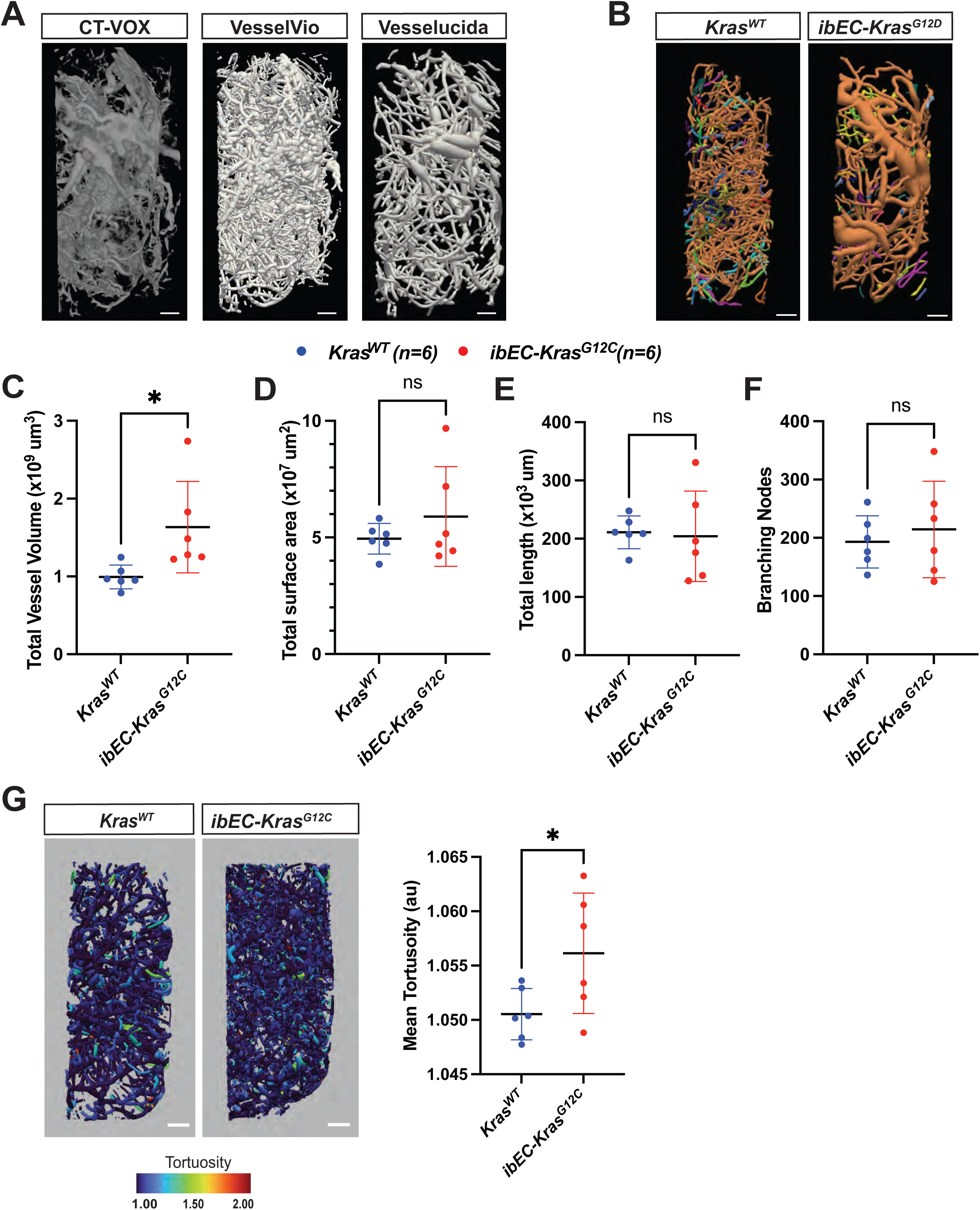
i*b*EC*-Kras^G12C^*–induced BAVMs feature extensively remodeled vessels. (**A**) Isolation of a region of interest [ROI] containing a bAVM, as opposed to whole brain reconstructions and analysis from the entire mouse brain as seen in Supplemental Figure 6, after micro-CT imaging and NRECON reconstruction followed by 3D rendering using CT-VOX, CT-AN, VesselVio, and Vesselucida. (**B**) Vessel tracing of the ROI that usually contains a bAVM (Bregma -3 to -5) in *ibEC-Kras^G12C^* mutants. Vessels are color coded to denote directly-connected vessel networks. Scale bar = 500 μm. (**C**) Quantification of vessel volume within the ROI in control *Kras^WT^* and *ibEC-Kras^G12C^* mice (each dot indicates an ROI from a single animal). *p=0.0411 (student’s T-test). (**D**) Quantification of total surface area for all vessels within the ROI from control *Kras^WT^* and *ibEC-Kras^G12C^* mice (each dot indicates an ROI from a single animal). p=0.333 (student’s T-test). (**E**) Quantification of total vessel length in the ROI from control *Kras^WT^* and *ibEC-Kras^G12C^* mice (each dot indicates an ROI from a single animal). p=0.627 (student’s T-test). (**F**) Quantification of branching node number within the ROI from control *Kras^WT^* and *ibEC-Kras^G12C^* mice (each dot indicates an ROI from a single animal). p=0.651 (student’s T-test). (**G**) Renderings of vessel tortuosity, utilizing VesselVio, within the ROI from control *Kras^WT^* and *ibEC-Kras^G12C^* mice on the left and quantification on the right (each dot indicates an ROI from a single animal). Scale bar = 500 μm. *p= 0.0231 (student’s T-test).

**Supplemental Figure 11.**
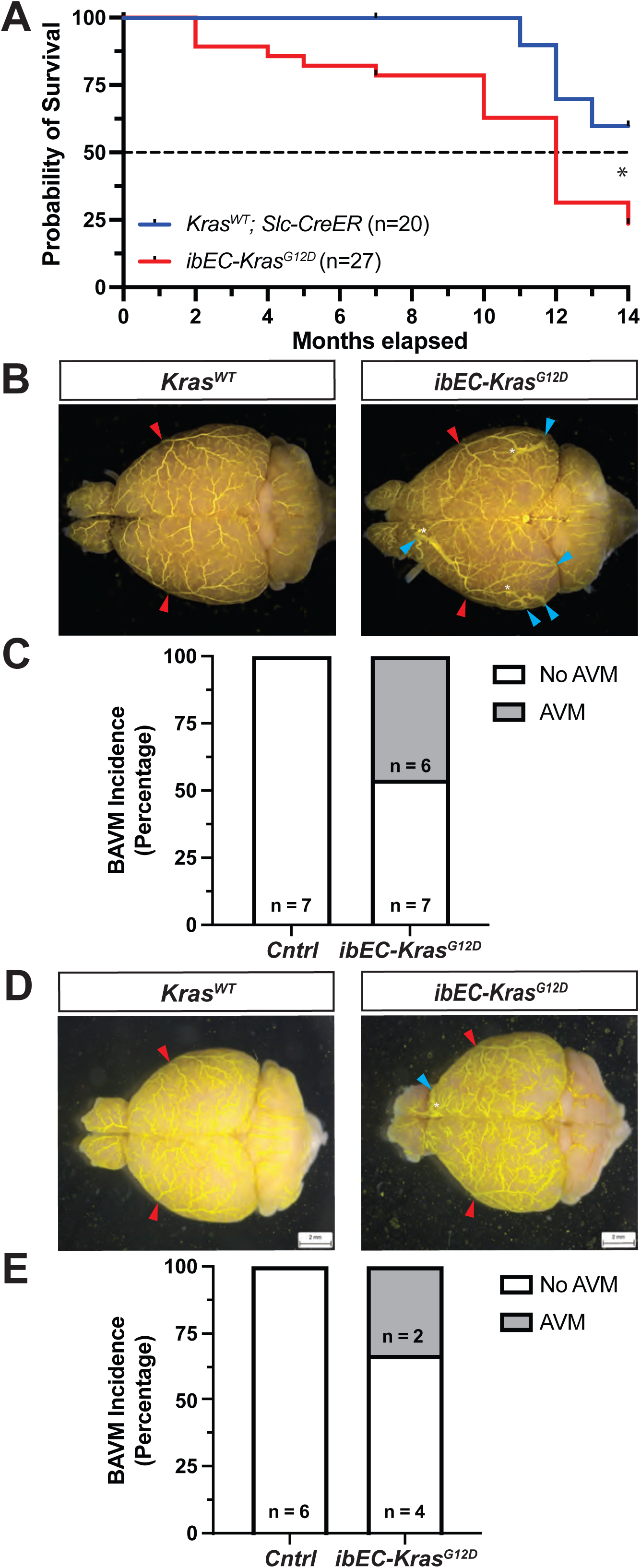
The survival rate of *ibEC-Kras^G12D^* mice decreases after 2 months of age. (**A**) While no differences in survival were evident between the two genotypes at 2 months, analysis over 14 months (Mantel-Haenszel, hazard ratio = 0.3064; Chi-square = 5.537, Gehan-Wilcoxon post-test, 1 degree of freedom, p=0.0186) demonstrates that *ibEC-Kras^G12D^* mice (n=27) are 3.263 times (inverse of the hazard ratio) more likely to die than *Kras^WT^* littermates (n=20). (**B**) Representative phase microscopy images of the dorsal surfaces of the brain after perfusions with Vascupaint contrast agent. Scale bar = 2000 μm. (**C**) Quantification of bAVM incidence, as determined by vascupaint perfusion, at 7 months of age after post tamoxifen induction at P1. (**D**) Representative phase microscopy images of the dorsal surface of the brain after perfusion with vascupaint. Scale bar = 2000 μm. (**E**) Quantification of bAVM incidence, as determined by vascupaint perfusion at 14 months of age after post tamoxifen induction at P1. Red carets denote the origin of the MCA on either the left or right side of the brain, respectively. Blue carets denote dilated draining veins, while the white asterisk denotes an arteriovenous fusion.

**Supplemental Figure 12.**
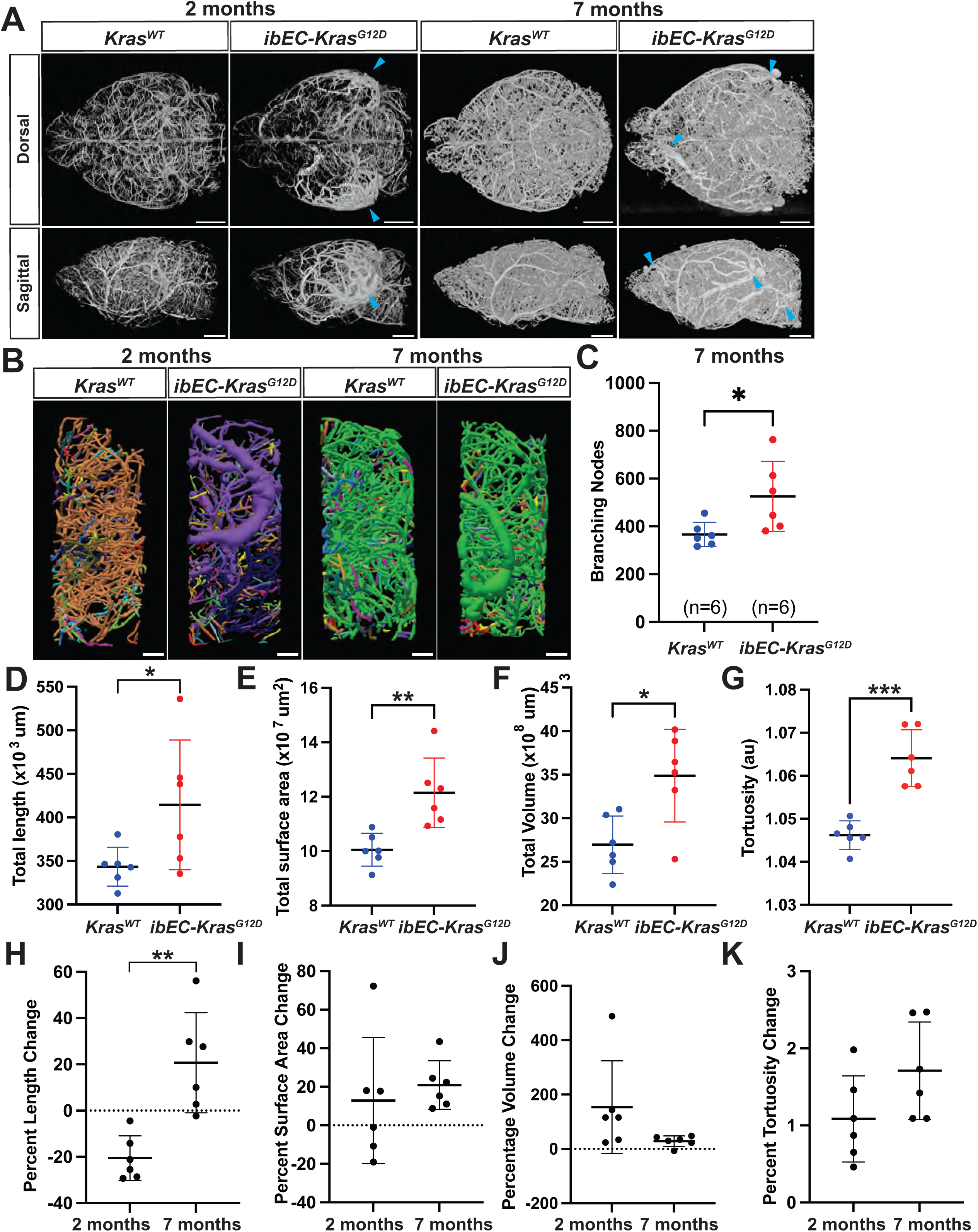
Aging does not induce remodeling in *ibEC-Kras^G12D^* mice. (**A**) Micro CT images following perfusion of the arterial circulation network at 2 months and 7 months of age for *Control* and *ibEC-Kras^G12D^* brains (top row = dorsal view; bottom row = sagittal view). Blue carets denote vascular anomalies. Scale bars = 2000 μm. (**B**) Vesselucida rendering of the bAVM ROI located in the -3 to -5 section from control *Kras^WT^* and *ibEC-Kras^G12D^* mice. Vessels are color coded to denote a directly-connected vascular network. Scale bars = 500 μm. (**C**) Quantification of the number of branching nodes in control *Kras^WT^* and *ibEC-Kras^G12D^* mice at 7 months of age (each dot indicates an ROI from a single animal). *p= 0.0308 (student’s T-test). (**D**) Quantification of vessel length in 7 month old control *Kras^WT^* and *ibEC-Kras^G12D^* mice from the -3 to -5 ROI. *p= 0.0491 (student’s T-test). (**E**) Quantification of total vessel surface area in 7 month old control *Kras^WT^* and *ibEC-Kras^G12D^* mice from the -3 to -5 ROI. **p=0.0045 (student’s T-test). (**F**) Quantification total vessel volume in 7 month old control *Kras^WT^* and *ibEC-Kras^G12D^* mice from the -3 to -5 ROI. *p= 0.0112 (student’s T-test). (**G**) Quantification of mean vessel segment tortuosity in 7 month old control *Kras^WT^* and *ibEC-Kras^G12D^* mice from the -3 to -5 ROI. ***p= 0.0001 (student’s T-test). (**H**) Percent change of vessel length. Each data point was calculated as the difference between the *ibEC-Kras^G12D^* ROI vessel length minus the average *Kras^WT^* ROI vessel length divided by the average *Kras^WT^* ROI vessel length, and multiplied by 100 % (Percent change = ((Mutant ROI vessel length – Average control ROI vessel length)/Average control ROI vessel length))*100. p=0.0017 (student’s T-test) (**I**) Percent change of the vessel surface. p=0.5869 (student’s T-test). (**J**) Percent change of the vessel volume. p=0.1056 (student’s T-test). (**K**) Percent change of the vessel tortuosity. p=0.0997 (student’s T-test).

**Supplemental Figure 13.**
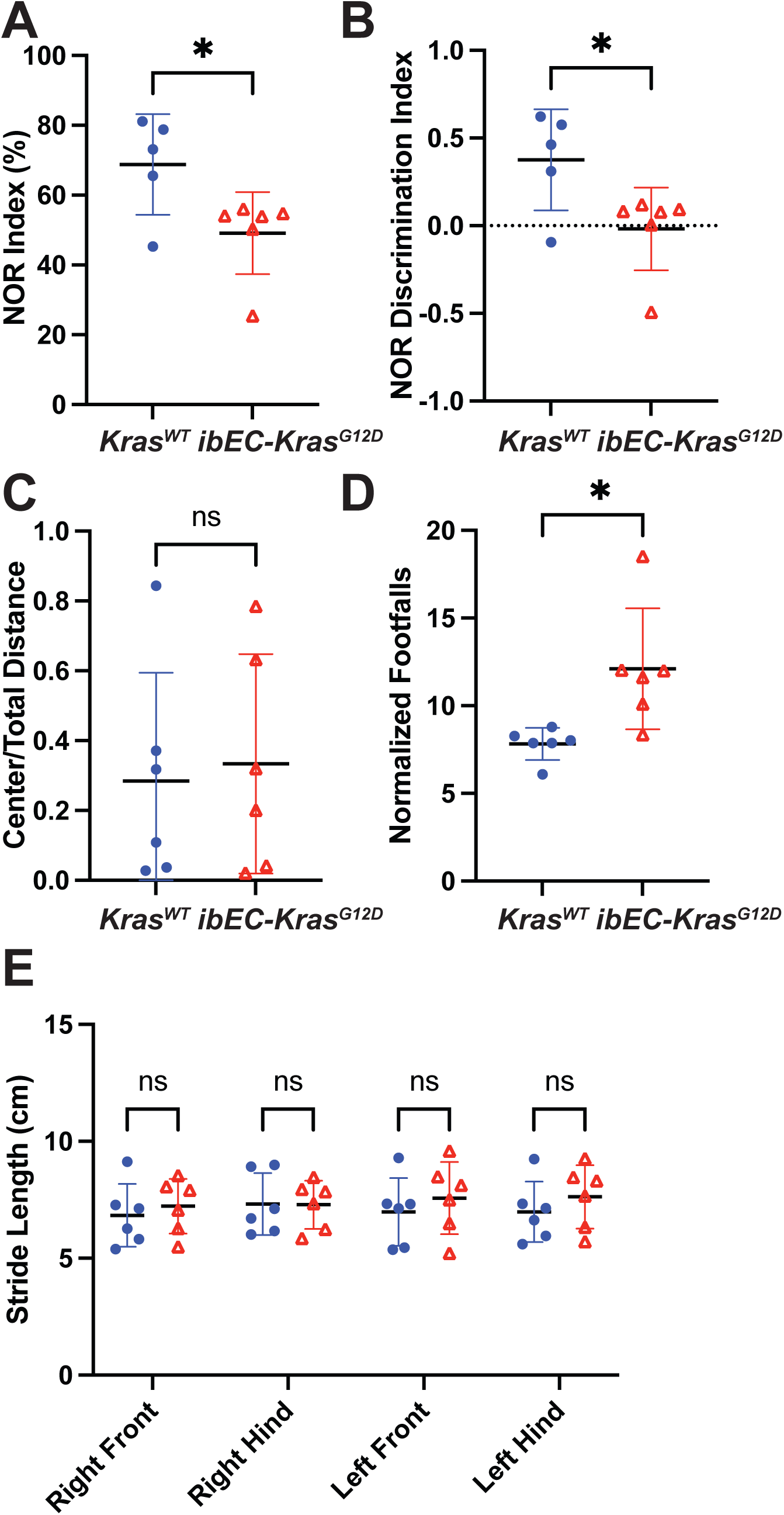
Locomotion and cognitive function are altered in mutant *ibEC-Kras^G12D^* mice at 7 months of age. (**A**) Evaluation of cognition using the novel object recognition (NOR) test, which is designed to asses short and long-term recognition memory, revealed a lower NOR index ((Time spent exploring novel object)/(Total time exploring both novel and familiar object)) in *ibEC-Kras^G12D^* mice than control littermates at 7 months of age. Each filled circle represents a mouse without a bAVM (as determined by micro CT analysis). Each triangle represents a mouse with a confirmed bAVM (*p=0.0339, student’s T-test). (**B**) The NOR discrimination index, or discrimination ratio, ((Time spent exploring novel object – Time spent exploring familiar object)/(Total time exploring both novel and familiar object)) is also different between *Kras^WT^* mice and *ibEC-Kras^G12D^* with confirmed bAVMs. *p=0.0340 (student’s T-test). (**C**) The open field activity test, which examines general activity and exploration and functions as a proxy for anxiety, revealed no change in the time spent in the center of the field compared to the total distance traveled between *Kras^WT^* mice vs *ibEC-Kras^G12D^* mice with confirmed bAVM (*p=0.9539, student’s T-test). (**D**) The parallel rod footfall (PRF) test, a simple measure of sensorimotor function and coordination in rodents, comparing *Kras^WT^* mice vs *ibEC-Kras^G12D^* with bAVMs (**p=0.0036, student’s T-test). (**E**) The CatWalk test, a system used for gait analysis in rodents shows that parameter stride length is indistinguishable, in any of the paws, between *Kras^WT^* vs *ibEC-Kras^G12D^* mice with AVMs. Student’s T-test were conducted for each comparison. For comparison at the right front p=0.6035; for right hindlimb, p=0.9624; for left front limb, p=0.5129; for left hindlimb, p=0.4204.

**Supplemental Figure 14.**
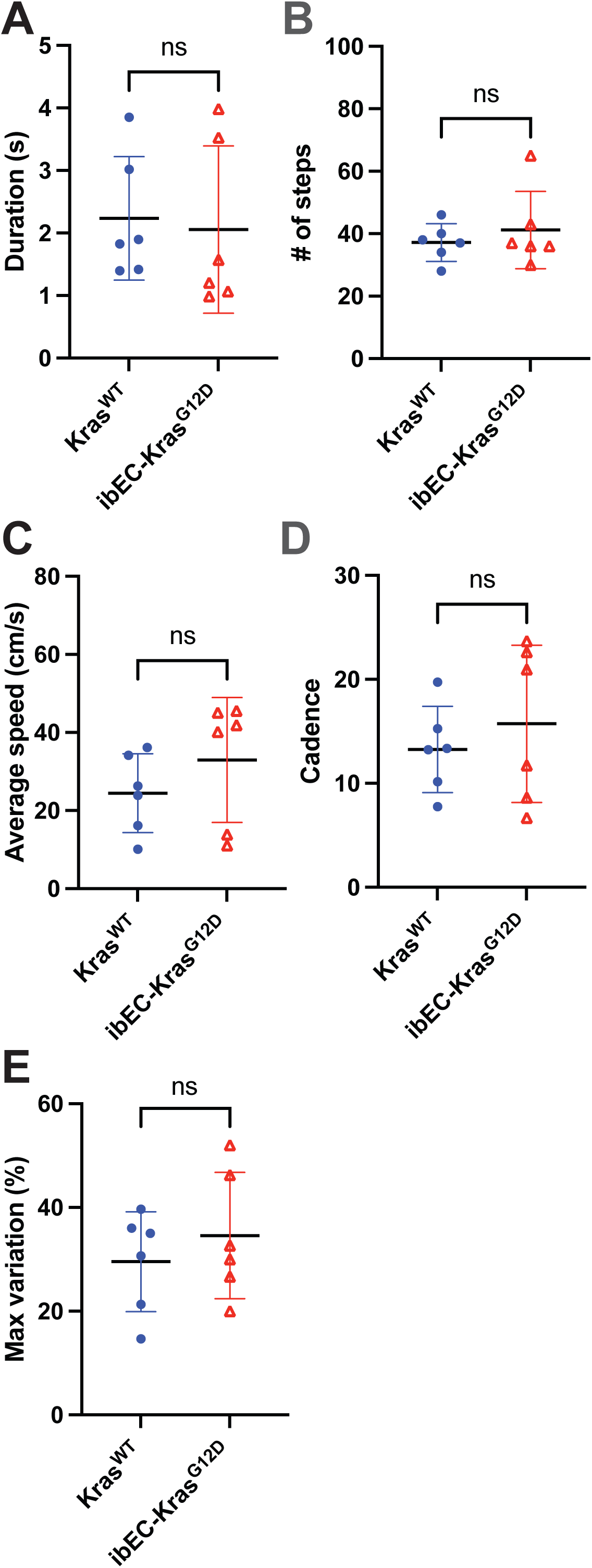
Gait analysis shows no locomotor deficits in *ibEC-Kras^G12D^* mice. (**A**) Duration of the run, in seconds. Each filled circle represents a mouse without a bAVM (as determined by micro-CT analysis). Each triangle represents a mouse with a confirmed bAVM. Duration of the run comparing *Kras^WT^* mice and only *ibEC-Kras^G12D^* with a confirmed bAVM. p=0.6473, student’s T-test. (**B**) Number of steps taken between *Kras^WT^* mice and *ibEC-Kras^G12D^* with confirmed bAVMs. p=0.4678, student’s T-test. (**C**) Average speed during the run between *Kras^WT^* mice and *ibEC-Kras^G12D^* with confirmed bAVMs. p=0.0643, student’s T-test. (**D**) Cadence between *Kras^WT^* mice and *ibEC-Kras^G12D^* with confirmed bAVMs. p=0.5633, student’s T-test. (E) Maximal variation between *Kras^WT^* mice and *ibEC-Kras^G12D^* with confirmed bAVMs. p=0.4445, student’s T-test.

**Supplemental Figure 15.**
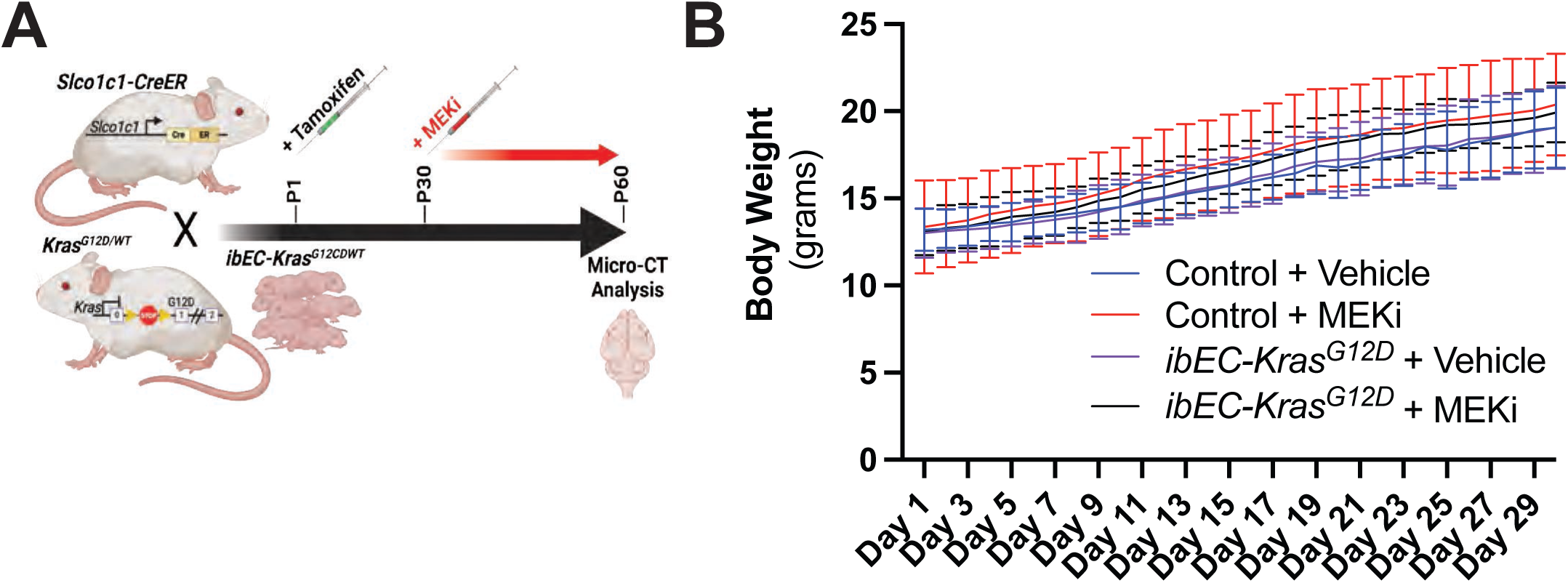
MEKi treatment does not impact weight gain or viability in mice over the treatment window. **(A)** Schematic for the generation of *ibEC-KrasG12D* experimental mice and the drug treatment window from P30 to P60. (**B**) Graphical representation of the animals’ weight over the treatment window shows that MEKi treatment had no significant effect on weight gain compared to treatment with the vehicle control.

**Supplemental Figure 16.**
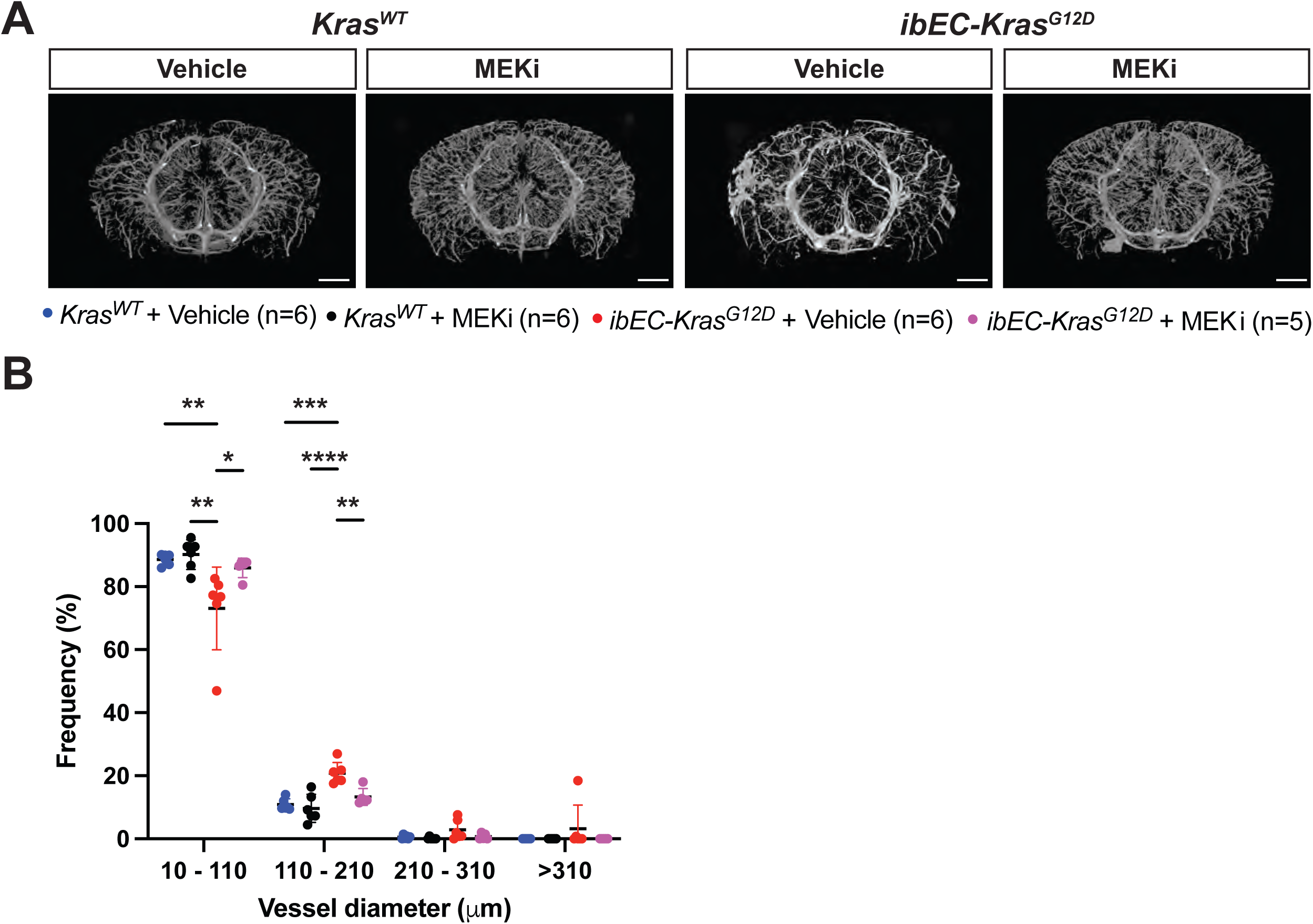
Treatment with a MEK inhibitor normalizes mutant KRAS-induced changes in the posterior cerebrovasculature. (**A**) Vessel tracing of the ROI that contains a bAVM (Bregma -3 to -5) in *ibEC-Kras^G12D^* mutants. All groups included: *ibEC-Kras^G12D^* mice and *Kras^WT^* mice treated with vehicle or MEK inhibitor. Vessels are color-coded to denote directly-connected vessel networks. Scale bar = 500 um. (**B**) Quantification of vessel volume within the ROI in control *Kras^WT^* and *ibEC-Kras^G12D^* mice (each dot indicates an ROI from a single animal) revealed a non-significant decrease in total vessel volume in *ibEC-Kras^G12D^* mice treated with MEK inhibitor compared to *ibEC-Kras^G12D^* mice treated with vehicle (p=0.0586). Similarly, total vessel volume is decreased in *Kras^WT^* mice treated with vehicle (p=0.0220) or MEKi (p=0.0246). (**C**) Quantification of total vessel surface area within the ROI from control *Kras^WT^* and *ibEC-Kras^G12D^* mice (each dot indicates an ROI from a single animal) treated with vehicle or MEK inhibitor. No significant change was observed. (**D**) Quantification of total vessel length within the ROI from control *Kras^WT^* and *ibEC-Kras^G12D^* mice (each dot indicates an ROI from a single animal). We observed an increase in total vessel length in *ibEC-Kras^G12D^* mice treated with MEK inhibitor compared to *ibEC-Kras^G12D^* mice treated with vehicle only (p=0.0012). Similarly, total vessel length was increased in *Kras^WT^* treated with vehicle (no significant, p=0.1731) or MEKi (p=0.0034) (**E**) Quantification of branching node number within the ROI from control *Kras^WT^* and *ibEC-Kras^G12D^* mice (each dot indicates an ROI from a single animal). We observed an increase of branching nodes in *ibEC-Kras^G12D^* mice treated with MEK inhibitor compared to *ibEC-Kras^G12D^* mice treated with vehicle only, but it was not significant (p=0.1164). Similarly, this was increased in *Kras^WT^* treated with vehicle (no significant, p=0.0584) or MEKi (p=0.0176) (**F**) Quantification of tortuosity within the ROI from *Kras^WT^* and *ibEC-Kras^G12D^* mice (each dot indicates an ROI from a single animal). We observed a decrease of tortuosity in *ibEC-Kras^G12D^* mice treated with MEK inhibitor compared to *ibEC-Kras^G12D^* mice treated with vehicle only, but it was not significant (p=0.1323). Similar decrease was observed in *Kras^WT^* treated with vehicle (p=0.0004) and MEKi (p=0.0002). One-way ANOVA was performed for each parameter followed by a Tukey’s multiple comparison test.

**Supplemental Figure 17.**
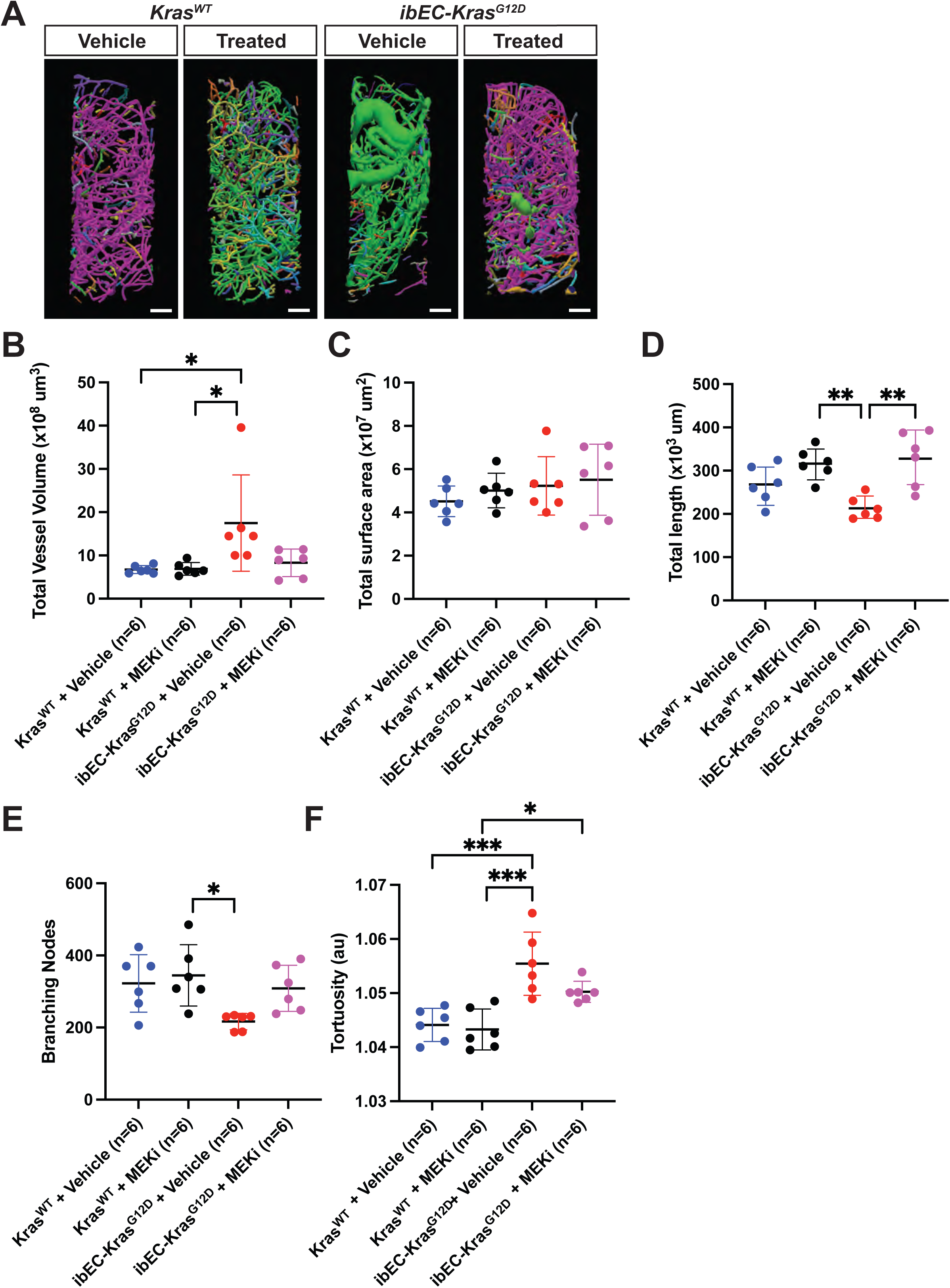
Vascular parameters within the ROI of *ibEC-Kras^G12D^* treated with MEK inhibitors normalize to those of *Kras^WT^* mice. (**A**) Vessel tracing of the ROI that contains a bAVM (Bregma -3 to -5) in *ibEC-Kras^G12D^* mutants. All groups included: *ibEC-Kras^G12D^* mice and *Kras^WT^* mice treated with vehicle or MEK inhibitor. Vessels are color-coded to denote directly-connected vessel networks. Scale bar = 500 μm. (**B**) Quantification of vessel volume within the ROI in control *Kras^WT^* and *ibEC-Kras^G12D^* mice (each dot indicates an ROI from a single animal). A non-significant decrease in total vessel volume was observed in *ibEC-Kras^G12D^* mice treated with MEK inhibitor compared to *ibEC-Kras^G12D^* mice treated with vehicle only, but it is not significant (p=0.0586). Similarly, total vessel volume is decreased in *Kras^WT^* mice treated with vehicle (p=0.0220) or MEKi (p=0.0246). (**C**) Quantification of total vessel surface area within the ROI from control *Kras^WT^* and *ibEC-Kras^G12D^* mice (each dot indicates an ROI from a single animal) treated with vehicle or MEK inhibitor. No significant change was observed. (**D**) Quantification of total vessel length within the ROI from control *Kras^WT^* and *ibEC-Kras^G12D^*mice (each dot indicates an ROI from a single animal). We observed an increase in total vessel length in *ibEC-Kras^G12D^* mice treated with MEK inhibitor compared to *ibEC-Kras^G12D^* mice treated with vehicle only (p=0.0012). Similarly, total vessel length was increased in *Kras^WT^* treated with vehicle (no significant, p=0.1731) or MEKi (p=0.0034) (**E**) Quantification of branching node number within the ROI from control *Kras^WT^* and *ibEC-Kras^G12D^* mice (each dot indicates an ROI from a single animal). We observed an increase of branching nodes in *ibEC-Kras^G12D^* mice treated with MEK inhibitor compared to *ibEC-Kras^G12D^* mice treated with vehicle only, but it was not significant (p=0.1164). Similarly, this was increased in *Kras^WT^* treated with vehicle (no significant, p=0.0584) or MEKi (p=0.0176) (**F**) Quantification of tortuosity within the ROI from control *Kras^WT^* and *ibEC-Kras^G12D^* mice (each dot indicates an ROI from a single animal). We observed a decrease of tortuosity in *ibEC-Kras^G12D^* mice treated with MEK inhibitor compared to *ibEC-Kras^G12D^* mice treated with vehicle only, but it was not significant (p=0.1323). Similar decrease was observed in control *Kras^WT^* treated with vehicle (p=0.0004) and MEKi (p=0.0002). One-way ANOVA was performed for each parameter followed by a Tukey’s multiple comparison test.

## Supplemental Videos

**Supplemental Video 1. Micro-CT 3D scan and virtual sections show *Kras^WT^* brains feature normal cerebrovasculature.** A representative scan from a 2 month old, control *Kras^WT^* brain from the *ibEC-Kras^G12D^* experimental cohort, created using CTVox software. The initial view is along a vertical axis, followed by virtual horizontal sections, followed by a horizontal rotation and then sectioned in the coronal plane. Note that no brain arteriovenous malformations are present in this sample.

**Supplemental Video 2. Micro-CT 3D scan and virtual sections show *ibEC-Kras^G12D^* brains feature bAVMs with dilated and tortuous vessels.** A representative scan from a 2 month old, *ibEC-Kras^G12D^* brain, created using CTVox software. The initial view is along a vertical axis, followed by virtual horizontal sections, followed by a horizontal rotation and then sectioned in the coronal plane. In the sample, bAVMs are present on the left and right hemispheres of the brain, respectively, located at Bregma -3 to -5 in the cortex.

**Supplemental Video 3. Micro-CT 3D scan and virtual sections show *Kras^WT^* brains feature normal cerebrovasculature.** A representative scan from a 2 month old, control *Kras^WT^* brain from the *ibEC-Kras^G12C^* experimental cohort, created using CTVox software. The initial view is along a vertical axis, followed by virtual horizontal sections, followed by a horizontal rotation and then sectioned in the coronal plane. Note that no brain arteriovenous malformations are present in this sample.

**Supplemental Video 4. Micro-CT 3D scan and virtual sections show *ibEC-Kras^G12C^* brains feature bAVMs with dilated and tortuous vessels.** A representative scan from a 2 month old, *ibEC-Kras^G12C^* brain, created using CTVox software. The initial view is along a vertical axis, followed by virtual horizontal sections, followed by a horizontal rotation and then sectioned in the coronal plane. In the video, five bAVMs are observed at Bregma +6 to +3 (1) in the left side of the olfactory bulbs. At Bregma -3 to -5 in the cortex, dorsal side (2), and in the left and right hemispheres (2).

